# Targeting Regnase-1 unleashes CAR T cell antitumor activity for osteosarcoma and creates a proinflammatory tumor microenvironment

**DOI:** 10.1101/2025.05.20.650777

**Authors:** Adeleye O. Adeshakin, Hao Shi, S. Scott Perry, Heather Sheppard, Phuong Nguyen, Xiang Sun, Peipei Zhou, Jean-Yves Métais, Trevor Cunningham, KC Anil, Liqing Tian, Vivek Peche, Mollie S. Prater, Deanna M. Langfitt, Shondra M. Pruett-Miller, Jason T. Yustein, Giedre Krenciute, Christopher DeRenzo, Hongbo Chi, Stephen Gottschalk

**Author notes:** These authors share senior authorship (Hongbo Chi, Stephen Gottschalk). Corresponding Authors Stephen Gottschalk, Department of Bone Marrow Transplantation and Cellular Therapy, St. Jude Children’s Research Hospital, 262 Danny Thomas Place, Memphis, TN 38105, Hongbo Chi, Department of Immunology, St. Jude Children’s Research Hospital, 262 Danny Thomas Place, Memphis, TN 38105.

## Abstract

Negative regulators of T cell function represent promising targets to enhance the intrinsic antitumor activity of CAR T cells against solid tumors. However, the endogenous immune ecosystem in solid tumors often represents an immunosuppressive therapeutic barrier to CAR T cell therapy, and it is currently unknown whether deletion of negative regulators in CAR T cells reshapes the endogenous immune landscape. To address this knowledge gap, we developed CAR T cells targeting B7-H3 in immune-competent osteosarcoma models and evaluated the intrinsic and extrinsic effects of deleting a potent negative regulator called Regnase-1 (Reg-1). Deletion of Reg-1 not only improved the effector function of B7-H3-CAR T cells but also endowed them with the ability to create a proinflammatory landscape characterized by an influx of IFNγ-producing endogenous T cells and NK cells and a reduction of inhibitory myeloid cells, including M2 macrophages. Thus, deleting negative regulators in CAR T cells enforces a non-cell-autonomous state by creating a proinflammatory tumor microenvironment.

## INTRODUCTION

Adoptive immunotherapy with chimeric antigen receptor-expressing T cells (CAR T cells) has been successful for CD19- or BCMA-positive hematological malignancies, leading to their FDA approval^1^. However, with few exceptions, CAR T cell therapy has been less effective in early phase clinical studies for solid tumors^2–9^. Second genetic modification of CAR T cells has evolved as a promising approach to augment CAR T cells, including transgenic expression of immunostimulatory molecules or the deletion of negative regulators^3,10–20^.

Among established negative regulators, Regnase-1 (Reg-1), a ribonuclease that is involved in regulation of immune responses^21–23^, has emerged as a promising candidate. We and other investigators have demonstrated that, upon Reg-1 knock out (KO), tumor-specific CD19-CAR T or T cell receptor (TCR)-transgenic CD8^+^ T cells persist longer, express more effector molecules, and have increased oxidative phosphorylation (OXPHOS) compared to unmodified T cells^10,11,24,25^. Mechanistic studies revealed that the basic leucine zipper transcription factor ATF-like (BATF) is the key molecule that mediates Reg-1 KO-mediated reprogramming of CD8^+^ T cells^10^.

While these studies highlight the benefit of Reg-1 KO on improving the function of adoptively transferred T cells, little is known about the efficacy of Reg-1 KO CAR T cells in immune-competent models for solid tumors, with only one study demonstrating the benefit of Reg-1 KO in murine anti-leukemia CD19-CAR T cells^11^. In addition, the effect of Reg-1 KO CAR T cells on endogenous immune cells has not been examined in any immune-competent tumor model. These represent critical knowledge gaps, as the endogenous immune landscape of solid tumors often contributes to reduced CAR T cell therapeutic efficacy^26^. Here, we evaluated the effects of Reg-1 KO on the cell-autonomous and non-cell-autonomous functions of CAR T cells targeting B7-H3 (CD276), an antigen expressed on a broad range of pediatric and adult solid tumors^27–31^, in immune-competent osteosarcoma (OS) models. Notably, B7-H3-CAR T cells show significant antitumor activity in preclinical models and early phase clinical testing demonstrates safety, albeit with limited efficacy^27,29,32–34^. We show that Reg-1 KO B7-H3-CAR T cells have improved effector function *in vivo* and create an immunostimulatory lung microenvironment. Specifically, comprehensive flow cytometric and single cell RNA-sequencing analyses revealed an influx of interferon (IFN)γ-producing CD4^+^ and CD8^+^ T cells, as well as NK cells, whereas there was a reduction of inhibitory myeloid cells, including M2 macrophages. Therefore, targeting Reg-1 has both cell-autonomous effects to improve the effector function of CAR T cells and non-cell-autonomous effects to enable them to create a proinflammatory microenvironment *in vivo*.

## RESULTS

### Reg-1 KO improves the antitumor activity of B7-H3-CAR T cells *in vivo*

To rigorously establish the therapeutic effects of targeting Reg-1 for CAR T cell therapy, we generated Reg-1 and control (Ctrl) KO B7-H3-CAR T cells, as well as Reg-1 KO CAR T cells targeting an irrelevant antigen (SP6). Naive CD8^+^ T cells derived from Cas9-transgenic mice (CD45.1^+^) were co-transduced with retroviral vectors encoding the respective CAR or single guide (sg) RNA linked to an ametrine reporter. Median double transduction efficacy was 42% (**Fig. 1A, Fig. S1A**), and Reg-1 gene editing efficiency was confirmed by indel analysis (**Fig. 1B, Fig. S1B,C**). To evaluate the specificity of B7-H3-CAR T cells, we used murine B7-H3-positive (F331, M1199) or B7-H3-negative (F331 B7-H3-KO) OS cell lines (**Fig. S2**). Reg-1 and Ctrl KO B7-H3-CAR T cells readily killed F331 and M1199 cell lines, in contrast to non-transduced (NT) or Reg-1 KO SP6-CAR T cells (**Fig. 1C, Fig. S3**). Likewise, Ctrl or Reg-1 KO B7-H3-CAR T cells showed no or minimal killing of F331 B7-H3 KO cells, respectively, verifying antigen specificity (**Fig. 1C, Fig. S3**). To examine the effects of Reg-1 KO on cytokine/chemokine production we analyzed CAR T cells post generation (**Fig. 1D**) and after coculture with F331 cells (**Fig. 1E**). Post generation, Reg-1 KO B7-H3-CAR T cells produced significantly more GM-CSF and IFNγ than Ctrl KO B7-H3-CAR T cells (**Fig. 1D**). After coculture with F331 cells, as compared to Reg-1 KO SP6-CAR T cells, Reg-1 and Ctrl KO B7-H3-CAR T cells produced more TH1 cytokines (GM-CSF, IFNγ, TNFα, IL-1α, IL-2, IL-3, IL-15), TH2 cytokines (IL-6, IL-10, IL13, LIF), and chemokines (IP10, LIX, MCP-1, MIP-1α, MIP-1β, MIG, RANTES), with no differences observed between Ctrl and Reg-1 KO CAR T cells (**Fig. 1E**). Reg-1 KO B7-H3-CAR T cells also produced more IL-7 and CXCL1 than Reg-1 KO SP6-CAR T cells (**Fig. 1E**). Thus, we have successfully developed Reg-1 KO B7-H3-CAR T cells with high specificity and efficacy.

**Figure 1:**
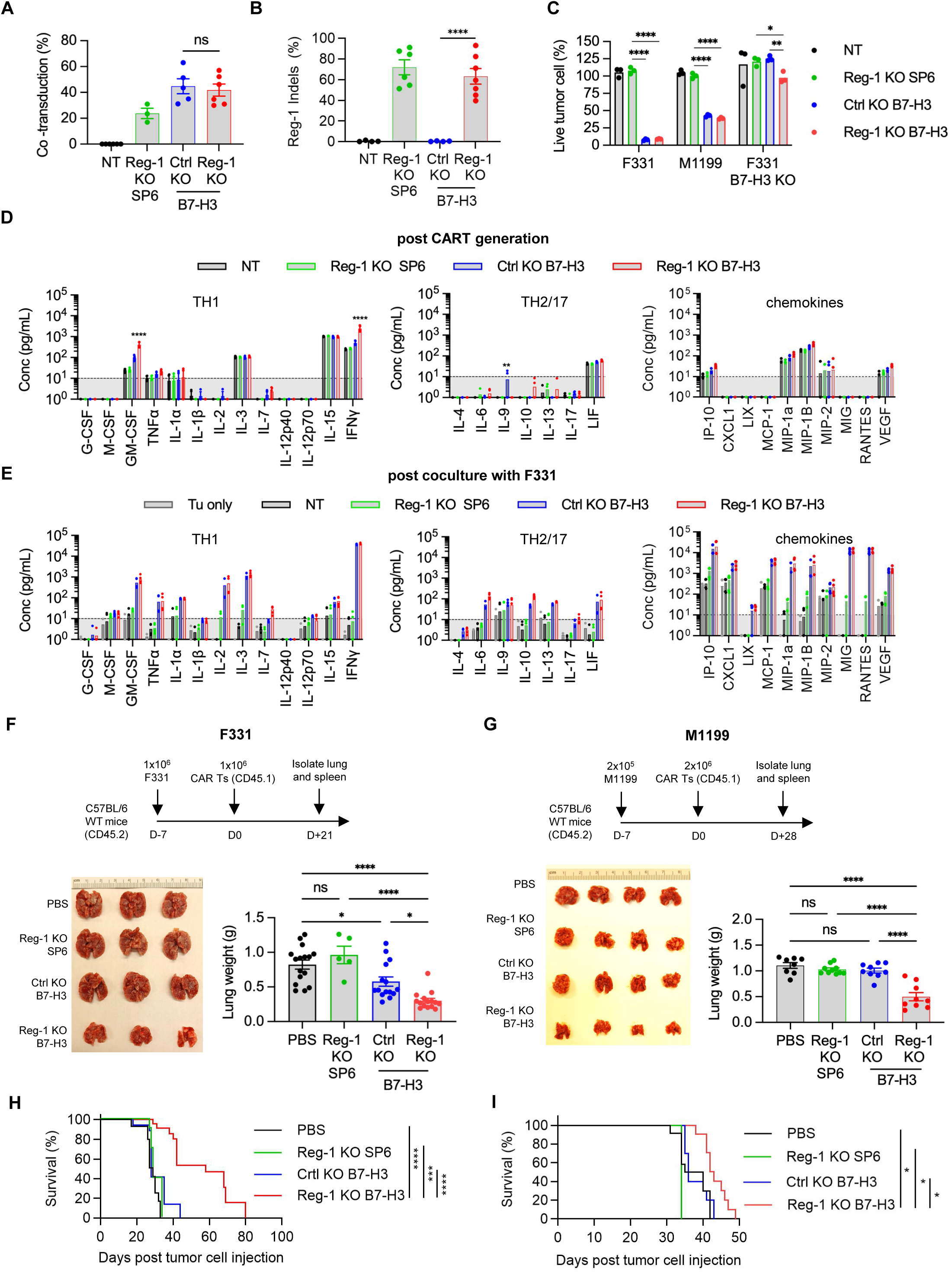
Reg-1 KO improves the antitumor activity of B7-H3-CAR T cells *in vivo*. (**A**) Co-transduction efficiency of indicated sgRNA (ametrine) and CAR in NT, (Ctrl) KO B7-H3-CAR, Reg-1 KO SP6-CAR, or Reg-1 KO B7-H3-CAR T cells evaluated by flow cytometry, n=3–6 pooled from multiple independent experiments (one-way ANOVA with Tukey’s test for multiple comparisons). (**B**) Insertion and deletion (Indel) analysis of the Reg-1 gene in indicated CAR T cells, n=4–7 pooled from multiple independent experiments (one-way ANOVA with Tukey’s test for multiple comparisons). (**C**) 3-(4,5-dimethylthiazol-2-yl)-5-(3-carboxymethoxyphenyl)-2-(4-sulfophenyl)-2H-tetrazolium (MTS) assay with B7-H3-positive (F331, M1199) and B7-H3-negative (B7-H3 KO F331) targets at an effector to target (E:T) ratio of 2:1; n=3 (two-way ANOVA with Tukey’s test for multiple comparisons). (**D,E**) Multiplex analysis of cytokines and chemokines post (**D**) CAR T-cell production (two-way ANOVA with Tukey’s test for multiple comparisons) and (**E**) 48 hours after stimulation with F331 cells at 1:1 E:T ratio. Tumor cell and NT T cell group, n=3; all other conditions, n=5. (**F–I**) C57BL/6 mice (CD45.2^+^) were injected with 1×10^6^ F331 or 2×10^5^ M1199 OS cells intravenously (i.v.) and on day 7 received a single i.v. dose of the indicated CAR T cell (Cas9^+^CD45.1^+^) populations or PBS. (**F,G**) Experimental schematic (upper), representative macroscopic images of lungs (lower left) and lungs weight (lower right) on (**F**) day 21 for F331 (n=3 for images) or (**G**) day 28 for M1199. n=4 per group for images and n=8–10 per group for lungs weight post-T cell infusion. (**H,I**) Kaplan-Meier survival curve (**H**) F331, log-rank Mantel-Cox test, n=8 for Reg-1 KO SP6; all other groups, n=13 and (**I**) M1199, log-rank Mantel-Cox test, n=5 for Reg-1 KO SP6; all other groups, n=10; two-way ANOVA with Tukey’s test for multiple comparisons. *:p<0.05, **:p<0.01, ***:p<0.001, ****:p<0.0001.

To evaluate the functional effects of Reg-1 KO on the antitumor activity of B7-H3-CAR T cells *in vivo*, we focused on models for OS tumor colonization in the lung, which is the primary site of metastasis and a predictor of poor prognosis^35–37^. Specifically, F331 or M1199 lung tumor-bearing congenic mice (CD45.2^+^) received a single i.v. injection of Reg-1 KO B7-H3-CAR, Ctrl KO B7-H3-CAR, Reg-1 KO SP6-CAR T cells, or PBS only (**Fig. 1F,G**). Infusion of Reg-1 KO B7-H3-CAR T cells had no overt side effects in mice, with no weight loss observed (**Fig. S4**). Reg-1 KO B7-H3-CAR T cells had superior antitumor activity compared to all other experimental groups, as revealed by lung weight comparison, and visual and histopathological observations, on day 21 (F331) or day 28 (M1199) post infusion (**Fig. 1F,G, Extended Data Fig. 1**). This resulted in a significant survival advantage of mice treated with Reg-1 KO B7-H3-CAR T cells in both tumor models (**Fig. 1H,I**). IHC analysis revealed that F331 and M1199 tumors continued to express B7-H3 compared to F331 B7-H3 KO tumors (**Fig. S5A**). Moreover, after Reg-1 KO B7-H3-CAR T cell therapy, progressive tumors continued to express B7-H3, thereby excluding antigen loss as a mechanism of immune escape (**Fig. S5B**). We also observed improved antitumor activity of Reg-1 KO B7-H3-CAR T cells in a subcutaneous (s.c.) F331 model (**Extended Data Fig. 2A-C**). Therefore, Reg-1 KO improves the antitumor activity of B7-H3-CAR T cells in multiple tumor models *in vivo*.

### Reg-1 KO enhances B7-H3-CAR T cell expansion *in vivo*

To gain mechanistic insight into the functional state of Reg-1 KO B7-H3-CAR, Ctrl KO B7-H3-CAR, or Reg-1 KO SP6-CAR T cells and their effects on endogenous immune cells, we euthanized F331 tumor-bearing mice on day 7 or day 21 or M119 tumor-bearing mice on day 7 or day 28 post CAR T cell infusion. We then performed phenotypic analysis of immune cells from lungs and spleens. In addition, blood samples were collected for complete blood counts (CBCs) on days 7, 14, and 21 (for both F331 and M119 models) and for cytokine/chemokine analysis on day 7 (only for F331 model) post CAR T cell infusion. Flow cytometric analysis of single-cell suspensions from lungs and spleens revealed a significantly higher frequency of Reg-1 KO B7-H3-CAR T cells compared to Ctrl KO B7-H3-CAR or Reg-1 KO SP6-CAR T cells on day 7 in both F331 and M119 tumor-bearing mice (**Fig. 2A,B**), which all declined to baseline levels by day 21/28 (**Fig. 2C,D**). An increased frequency of Reg-1 KO B7-H3-CAR T cells in tumors was confirmed by ISH (*in situ* hybridization) (**Extended Data Fig. 3**). At peak expansion (day 7), phenotypic analysis of Reg-1 KO B7-H3-CAR versus Ctrl KO B7-H3-CAR T cells revealed enhanced differentiation to an effector memory (EM; CD62L^-^CD44^+^) phenotype, with more pronounced effects observed in the F331 model (**Fig. 2E,F**). No differences in PD-1-expressing CAR T cells were observed between Ctrl and Reg-1 KO B7-H3-CAR T cells (**Fig. S6A,B**), suggesting selective effects of targeting Reg-1 on CAR T-cell state.

**Figure 2:**
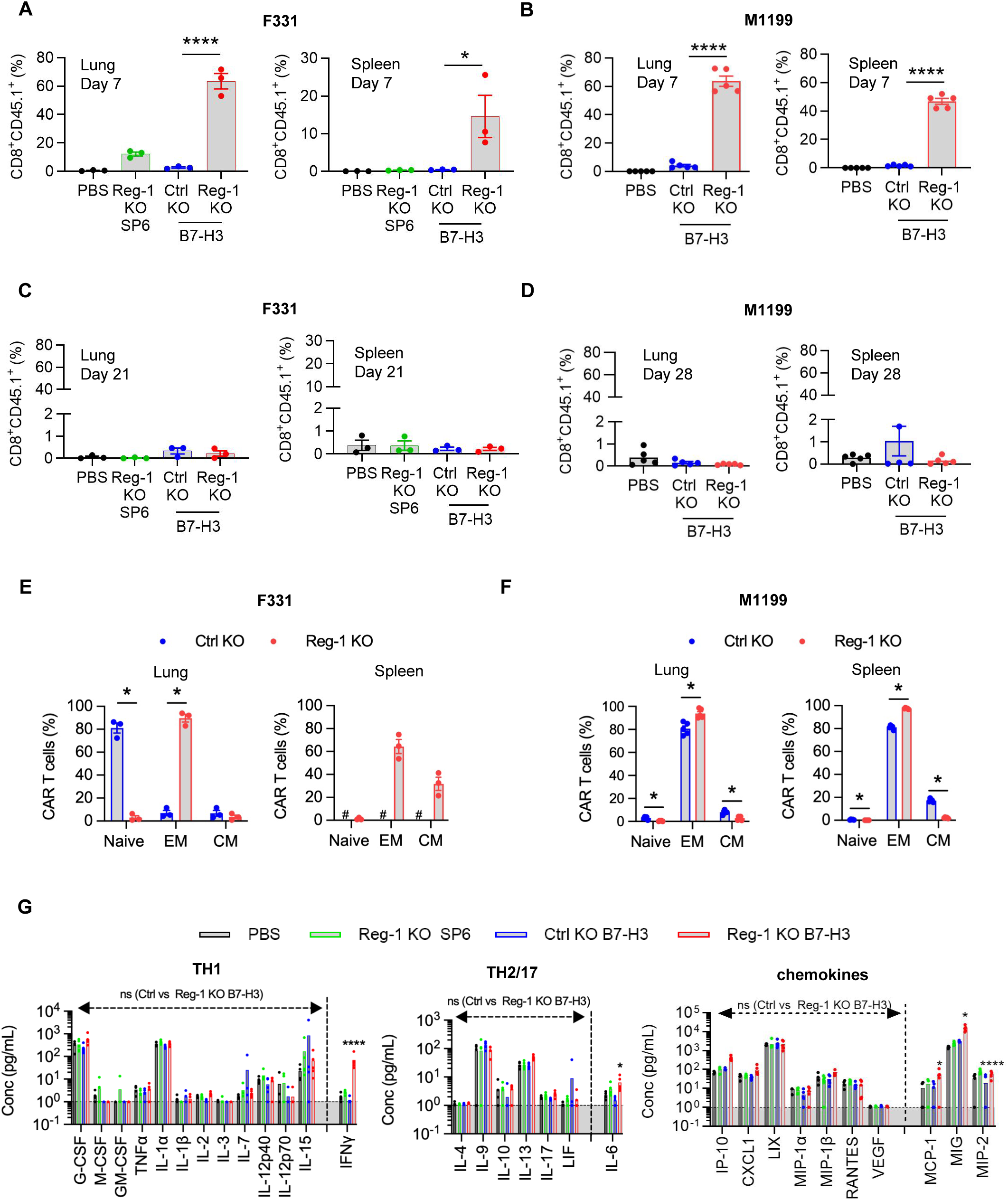
Reg-1 KO enhances B7-H3-CAR T cell expansion *in vivo*. C57BL/6 mice were injected with 1×10^6^ F331 or 2×10^5^ M1199 OS cells (i.v.) and on day 7 received a single i.v. dose of the indicated CAR T cell population or PBS. (**A–D**) Quantification of lungs- and spleens-infiltrating CD8^+^CD45.1^+^ T cells on day 7 for (**A**) F331 or (**B**) M1199 and (**C**) day 21 for F331 or (**D**) day 28 for M1119 post-T cell injection, as evaluated by flow cytometry. F331: n=3 per time point and M1199: n=5 per time point (one-way ANOVA with Tukey’s test for multiple comparisons). (**E,F**) Phenotyping of lungs- and spleens-infiltrating CAR T cell differentiation states, classified as naïve (CD62L^+^CD44^−^), effector memory (EM; CD62L^−^CD44^+^) and central memory (CM; CD62L^+^CD44^+^) subsets, in (**E**) F331 tumor-bearing mice (comparison could not be performed between Ctrl KO and Reg-1 KO due to insufficient Ctrl KO CAR T cells detected in the spleens; #: not done) and (**F**) M1199 tumor-bearing mice on day 7 post CAR T cell infusion (F331; n=3, M1199: n=5 unpaired student *t test*). (**G**) Multiplex analysis of cytokines in the serum on day 7 post treatment with indicated CAR T cells or PBS in the F331 model, n=5 per group (two-way ANOVA with Tukey’s test for multiple comparisons); *:p<0.05; **:p<0.01; ***:p<0.001; ****:p<0.0001.

We found that expansion of Reg-1 KO B7-H3-CAR T cells coincided with significant splenomegaly on day 7, which resolved by day 21/28 (**Extended Data Fig. 4A,B**). CBCs and cytokine/chemokine analyses revealed transient lymphopenia and monocytopenia (only F331 model) on day 7 (**Extended Data Fig. 4C,D**) and a significant fold-increase of IFNγ, IL-6, MCP-1, MIG, and MIP-2 in mice that received Reg-1 KO B7-H3-CAR T cells (**Fig. 2G**). Reg-1 KO B7-H3-CAR T cells caused distinct changes in the composition of the lung immune cell infiltrates, including a higher percentage of NK cells and a lower percentage of B cells (in M1199 model), with such effects not observed in the spleen (**Extended Data Fig. 5A,B**). Changes in the myeloid compartment (neutrophils, monocytes, M1 and M2 macrophages, and dendritic cells (DCs)) were model-dependent. For instance, lungs from F331 mice had increased neutrophils and reduced M2 macrophages following infusion of Reg-1 KO B7-H3-CAR T cells compared to Ctrl KO B7-H3-CAR T cells, while in the M1199 model no difference in neutrophils but increased M2 macrophages were observed (**Extended Data Fig. 5A,B**). Among endogenous T cell populations, we did not observe consistent changes between the F331 and M1199 models regarding the percentages of CD3^+^, CD4^+^, and CD8^+^ T cells (**Extended Data Fig. 5A,B**) or various CD4^+^ T cell subsets (naive, EM, central memory (CM)) (**Fig. S7A,B**). Within the endogenous CD8^+^ T cells, Reg-1 KO B7-H3-CAR T cells increased EM and decreased naive subsets in both the lungs and spleens from F331 mice (**Fig. S7A**), whereas no significant differences were observed in the M1199 model (**Fig. S7B**). Finally, Reg-1 KO B7-H3-CAR T cells induced expression of PD-1- and PD-L1 in endogenous T cells and myeloid cells, respectively (**Extended Data Fig. 6A,B**).

In summary, Reg-1 KO induced antigen-dependent expansion and effector differentiation of B7-H3-CAR T cells, and created a systemic and pulmonary-specific proinflammatory environment, as revealed by alterations of the endogenous immune cell infiltrates. While the increased percentage of myeloid cells expressing PD-L1 raised the possibility of reprogramming of the myeloid compartment to an inhibitory state, combining Reg-1 KO B7-H3-CAR T cells with anti-PD-L1 treatment did not further improve their antitumor activity (**Fig. S8A,B**), highlighting the need for more detailed analyses, as conducted in the following experiments.

### Reg-1 KO enhances proinflammatory and metabolic signaling pathways in B7-H3-CAR T cells at the expansion phase

To determine the cellular and molecular mechanisms of Reg-1 function in CAR T cells, we compared the transcriptomics profiles between Ctrl KO and Reg-1 KO B7-H3-CAR T cells by performing scRNA-seq of CAR T cells isolated from the lungs and spleens of F331 tumor-bearing mice on day 5 post CAR T-cell infusion (**Fig. 3A**). Uniform manifold approximation and projection (UMAP) analysis revealed distinct clustering of Ctrl KO and Reg-1 KO B7-H3-CAR T cells in lungs and spleens, and an increased number of Reg-1 KO relative to Ctrl KO B7-H3-CAR T cells in both tissues (**Fig. S9A–E**). In addition, scTCR-seq analysis revealed that CAR T cells were more clonally expanded compared to non-CAR T cells (**Fig. S9F**). Reg-1 KO B7-H3-CAR T cells in the lungs expressed higher levels of multiple effector genes, including *Itgae* (encoding CD103)*, Ccr8, Bhlhe40, Pdcd1, Ifng,* and *Ifngr1,* and gene set enrichment analysis (GSEA) revealed significant enrichment of Hallmark gene signatures associated with TNFα signaling, IL-2-STAT5 signaling, and inflammatory responses, as compared to Reg-1 KO B7-H3-CAR T cells from the spleens (**Fig. 3B,C**). These results suggest preferential activation of Reg-1 KO B7-H3-CAR T cells by the metastatic tumors in the lung compared to the spleen.

**Figure 3:**
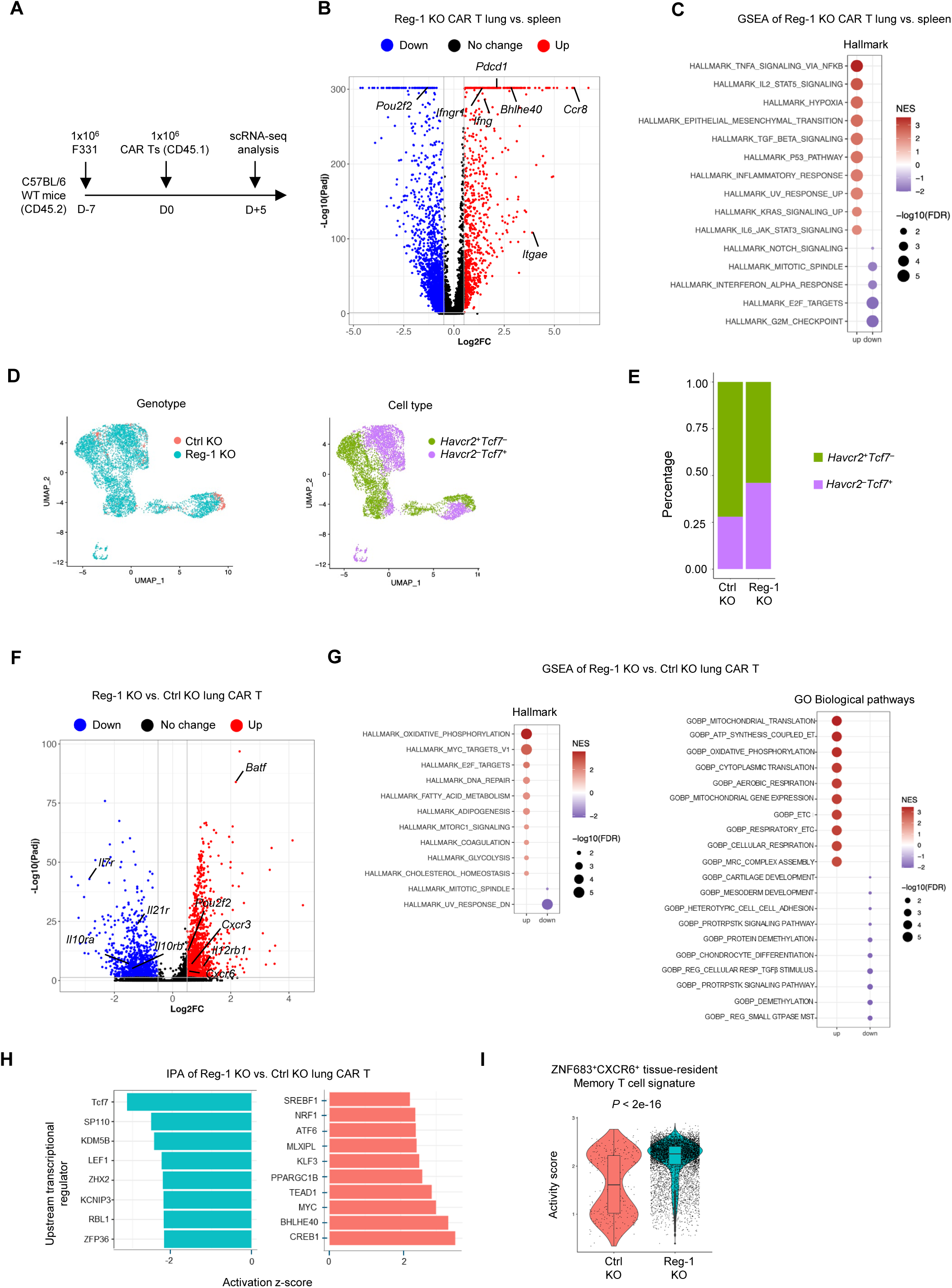
Reg-1 KO enhances pro-inflammatory and metabolic signaling pathways during the expansion phase of B7-H3 CAR T cells *in vivo*. (**A**) Experimental schematic: Mice received a single dose of 5×10^6^ Reg-1 KO or Ctrl KO B7-H3-CAR T cells (Cas9^+^CD45.1^+^) on day 7 post F331 tumor cell injection. Lungs and spleens were isolated on day 5 post CAR T cell infusion, and cells from 3 mice per condition were pooled for scRNA-seq analysis. (**B**) Volcano plot showing differentially expressed genes (Log2FC > 0.5 or < −0.5 and adjusted *P* < 0.05) in Reg-1 KO B7-H3-CAR T cells from the lungs versus spleens. (**C**) Top 10 significantly (FDR < 0.05) upregulated and top 5 downregulated pathways in Reg-1 KO B7-H3-CAR T cells from the lungs versus spleens, as revealed by gene set enrichment analysis (GSEA) using Hallmark gene sets. (**D**) Uniform Manifold Approximation and Projection (UMAP) of Reg-1 KO or Ctrl KO B7-H3 CAR T cells (left) and *Hvacr2* (encoding TIM-3) or *Tcf7* (encoding TCF1) expression (right) in the lungs. (**E**) Proportion of *Havcr2* and *Tcf7* in the Reg-1 KO and Ctrl KO B7-H3-CAR T cells in the lungs. (**F**) Upregulated (red) or downregulated (blue) genes (log2FC > 0.5 and adjusted *P* < 0.05) in Reg-1 KO B7-H3-CAR T cells compared to Ctrl KO B7-H3-CAR T cells. (**G**) Top 10 significantly (FDR < 0.05) upregulated and top 2 downregulated metabolism-related pathways (from Hallmark gene sets; left) or top 10 upregulated and downregulated GO biological pathways (right) in Reg-1 KO B7-H3-CAR T cells compared to Ctrl KO B7-H3-CAR T cells, as revealed by GSEA. (**H**) Ingenuity pathway analysis of transcriptional regulators of Reg-1 KO B7-H3-CAR T cells compared to Ctrl KO B7-H3-CAR T cells in the lungs. (**I**) Activity score of a CD8^+^ ZNF683^+^CXCR6^+^ tissue-resident memory T cell signature in Ctrl KO and Reg-1 KO B7-H3-CAR T cells in the lungs. ET: electron transport; ETC: electron transport chain; MRC: mitochondrial respiratory chain; PROTRPSTK: positive regulation of transmembrane receptor protein serine threonine kinase; REG: regulation; RESP: respiration; MST: mediated signal transduction.

We next compared the cellular differentiation states (based on *Havcr2* (encoding TIM-3) and *Tcf7* (encoding TCF1) expression) and gene expression profiles between Ctrl KO and Reg-1 KO B7-H3-CAR T cells collected from the lung. Compared to Ctrl KO B7-H3-CAR T cells, Reg-1 KO B7-H3-CAR T cells had a higher percentage of Havcr2^-^*Tcf7*^+^ and a lower percentage of *Havcr2^+^Tcf7^-^*cells (**Fig. 3D,E**). Differential gene expression analysis revealed that Reg-1 KO CAR T cells expressed higher levels of *Batf*, a transcription factor critical for effector differentiation of CD8^+^ T cells^10,38,39^. Cytokine receptor gene expression was either increased (*Il12rb1*) or decreased (*Il10ra*, *Il10rb*, *Il7r*, *Il21r*) in Reg-1 KO relative to Ctrl KO B7-H3-CAR T cells (**Fig. 3F**), highlighting that cytokine receptor expression is unlikely to predict T-cell functionality as observed in other models^13,40^. Reg-1 KO CAR T cells expressed higher levels of chemokine receptors (*Cxcr3*, *Cxcr6*) that orchestrate tumor-reactive T cell infiltration and function^41,42^. Based on GSEA, Reg-1 KO CAR T cells also upregulated cell proliferation signatures (Hallmark E2F targets) and metabolism-related signatures (Hallmark oxidative phosphorylation, Hallmark mTORC1, GOBP electron transport chain) compared to Ctrl KO CAR T cells (**Fig. 3G**). In addition, Ingenuity Pathway Analysis (IPA) revealed that Reg-1 KO CAR T cells upregulated activities for Bhlhe40, Myc, and PGC-1 (PPARGC1B), all of which are important for T cell mitochondrial function^43^ (**Fig. 3H**). Finally, Reg-1 KO B7H3-CAR T cells upregulated a pan-cancer ZNF683^+^CXCR6^+^ tissue-resident memory CD8^+^ T-cell signature^44^ (**Fig. 3I**), suggesting increased tissue residency features in the absence of Reg-1. These studies demonstrate that Reg-1 KO enhances proinflammatory and metabolic signaling pathways during the expansion phase of B7-H3-CAR T cells *in vivo*.

### Reg-1 KO B7-H3-CAR T cells reduce inhibitory myeloid cells and increase the frequency of IFNγ-producing T cells

Since our flow cytometric analyses on day 7 demonstrated that Reg-1 KO B7-H3-CAR T cells had significant effects on endogenous immune cells, we next determined whether these effects persisted at a later time point. To this end, F331 tumor-bearing mice were injected with PBS, Reg-1 KO B7-H3-CAR T cells, Ctrl KO B7-H3-CAR T cells or Reg-1 KO SP6-CAR T cells, followed by flow cytometric analysis of single-cell suspensions from lungs on day 21 post-CAR T-cell infusion. Two high-parameter flow cytometric experiments were conducted. The first profiled 19 unique cell markers (23 total fluorochromes; **Fig. 4, Fig. S10A, Fig. S11, Table S1, Table S2**), and the second 34 cell unique markers, respectively (38 total fluorochromes; **Fig. S10B, Fig. S12, Fig. S13, Tables S3, Table S4**). For all experimental groups, the two largest clusters consisted of myeloid and B cells, with T and NK cells being the next most abundant populations (**Fig. 4D, Fig. S13C**). CAR T cells were only detected after Reg-1 KO B7-H3-CAR T-cell infusion at low levels and co-expressed PD-1 and PD-L1 (**Extended Data** Fig. 7A,B). Within the myeloid compartment, there were a significantly higher frequencies of neutrophils and M1 macrophages, mirrored by a decrease in M2 macrophages defined by CD206 (MRC1) expression, upon Reg-1 KO B7-H3-CAR T-cell therapy (**Fig. 5A–C**).

**Figure 4:**
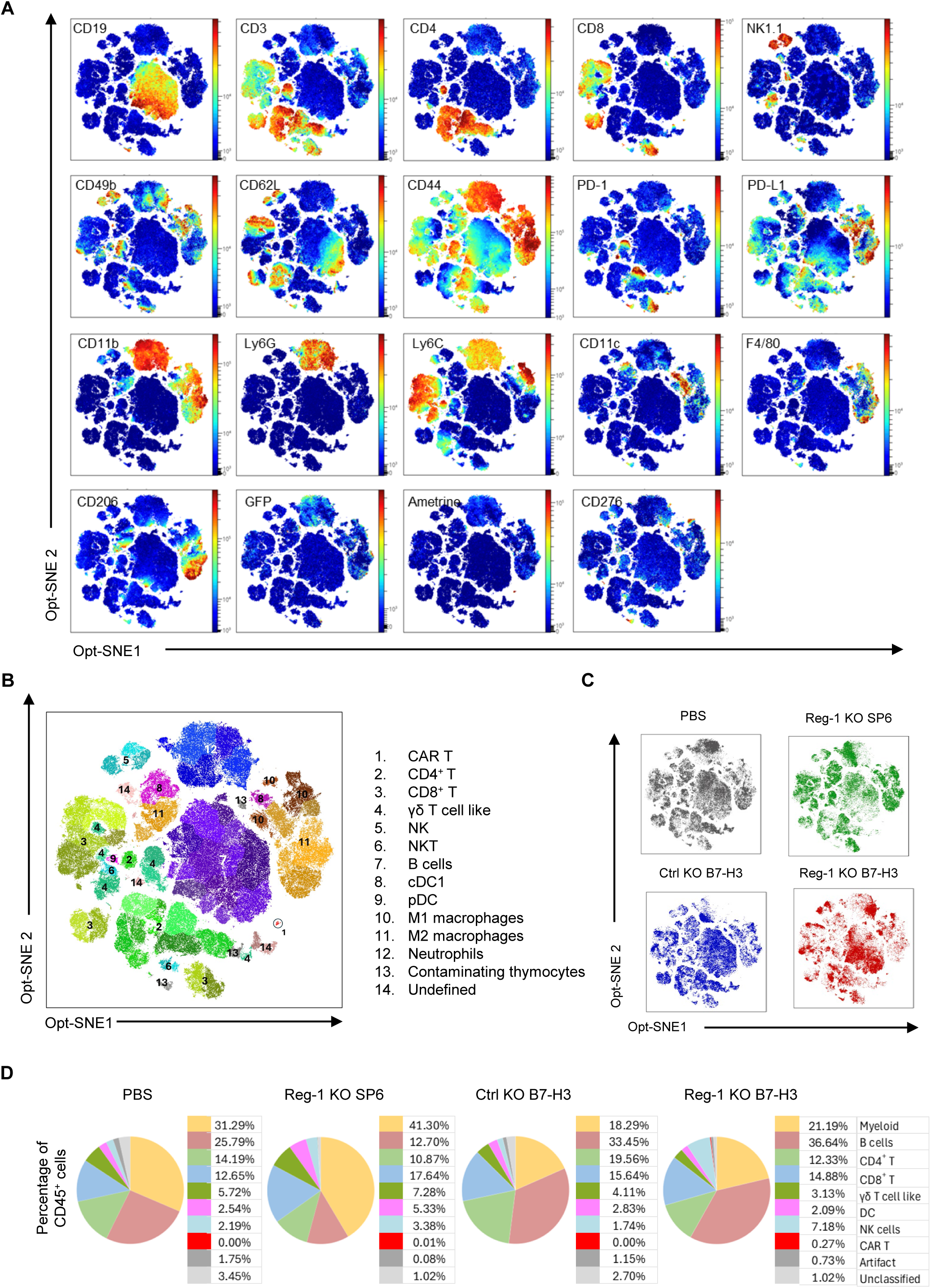
Global immune landscape changes post B7-H3-CAR T cell infusion. C57BL/6 mice (CD45.2^+^) were injected with 1×10^6^ F331 cells (i.v.) and on day 7 received a single i.v. dose of edited CAR T cells (Cas9^+^CD45.1^+^Ametrine^+^) population or PBS. On day 21 post CAR T cell infusion, mice were euthanized and lungs isolated for high dimensional flow cytometry analysis using a panel including 19 unique immune cell markers. An unbiased algorithm based opt-SNE analysis was performed on 26,000 viable CD45^+^ lung cell isolates per sample, followed by an unguided clustering in Flow-SOM utilizing elbow metaclustering. (**A**,**B**) Concatenated data from the lungs of 12 mice. (**A**) Heatmaps presenting the relative mean fluorescence intensity of the immune markers assayed, localized across the t-SNE clusters. (**B**) t-SNE cluster grouping according to immune subtypes based on root phenotypic marker expression as illustrated in **Fig S17** and detailed in **Table S2**. (**C**) t-SNE plots for indicated treatment conditions (n=3 mice per condition) displaying contributions of each test group to the concatenated data given in other panels, and illustrating distinct cluster aggregations between groups. (**D**) Fraction of CD45^+^ cells assigned to each immune cell subset, given as the mean from the 3 individuals in the indicated treatment group. t-SNE, t-distributed stochastic neighbor embedding. DC: dendritic cells, cDC1: conventional DC1, pDC: plasmocytoid DC.

**Figure 5:**
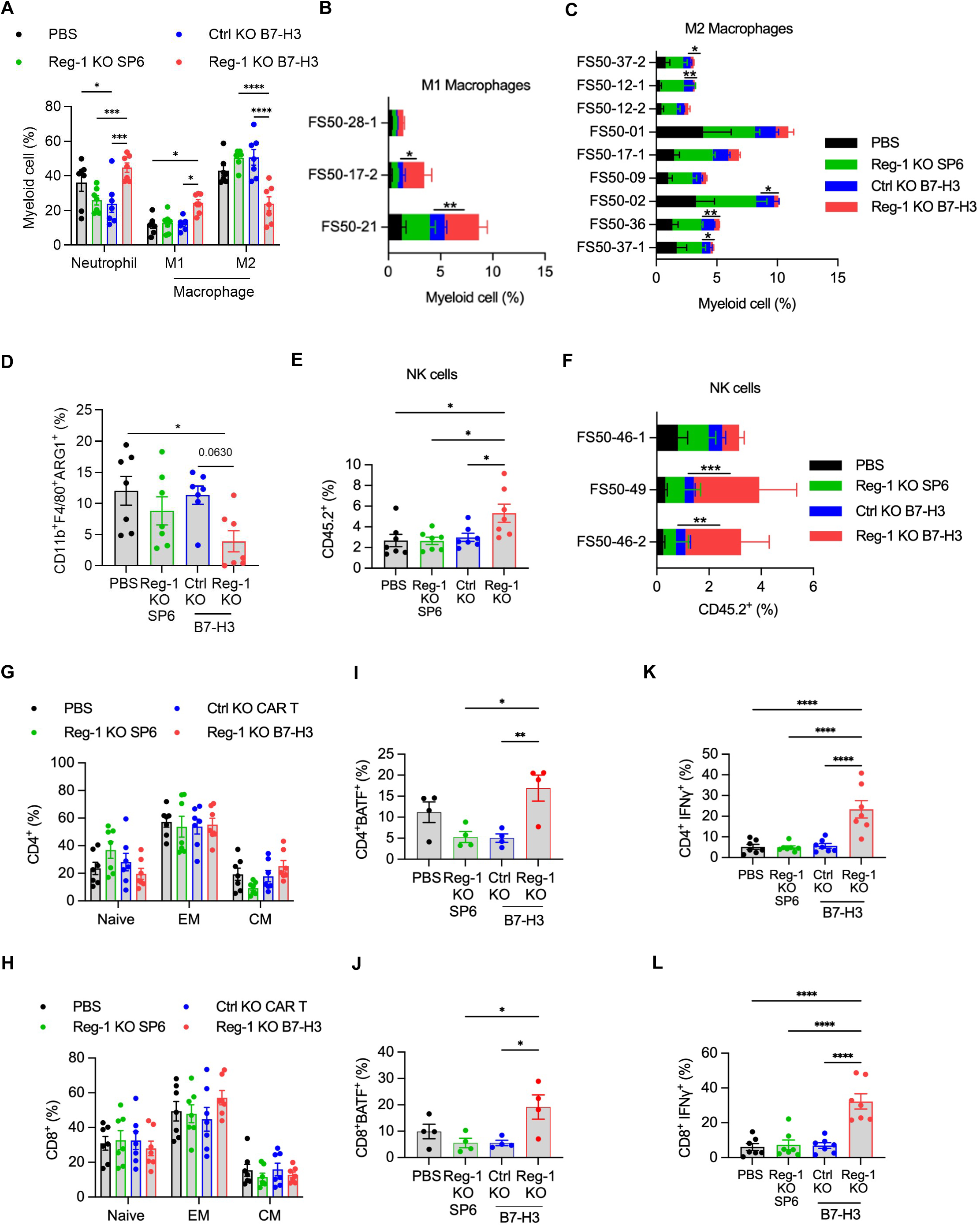
Reg-1 KO B7-H3-CAR T cells reduce endogenous inhibitory myeloid cells and increase pro-inflammatory immune cells. Extracellular marker expression data was pooled from the 2 high-parameter immunophenotyping experiments described for Fig. 4 and **Fig. S19**, totaling 7 lungs samples for each of the 4 indicated treatment conditions. Separate aliquots of lungs cell suspensions from all 28 mice treated in these studies were also probed with the intracellular flow cytometry panel (detailed in **Table S5**), as outlined in methods. (**A**) Percentages of indicated myeloid cell subsets, as identified based on the scheme outlined in **Fig. S17**. (**B**) Three M1 and (**C**) nine M2 macrophage clusters were identified, p <0.05 are shown for Ctrl KO versus B7-H3-CAR T cell comparison. (**D**) Frequencies of ARG1-expressing CD11b^+^ F4/80^+^ cells (total macrophages). (**E**) Percentages and (**F**) identified clusters of NK cells. In **F**, *p* values less than <0.05 are shown for Ctrl KO versus Reg-1 KO B7-H3-CAR T cell comparison. (**G–L**) Percentages of (**G,H**) naïve, EM, and CM populations, (**I,J**) BATF-expressing, or (**K,L**) IFNγ-expressing cells among CD4^+^ T cells (**G,I,K**) or CD8+ T cells (**H,J,L**), n=7 (**G,H,K,L**) or n = 4 (**I,J**) (two-way ANOVA with Tukey’s test for multiple comparisons) *:p<0.05; **:p<0.01; ***:p<0.001; ****:p<0.0001. EM: effector memory, CM: central memory. For panels **B, C**, and **F** data is shown for flow cytometric analysis (n=3).

We next analyzed lung single cell suspensions by flow cytometric analysis to profile expression of additional cell surface markers, intracellular cytokines and transcription factors (**Table S5**). Reg-1 KO B7-H3-CAR T cells induced a decrease in the frequency of CD11b^+^F4/80^+^ macrophages that also expressed arginase 1 (ARG1) (**Fig. 5D**), a hallmark of M2 macrophages that promotes T-cell dysfunction in the TME^45^. This finding was validated by IHC of lung tumors (**Extended Data** Fig. 8). There was also an increased frequency of NK cells (**Fig. 5E,F**). While no significant changes in the frequencies of endogenous naive, EM, and CM T cells were observed between treatment groups (**Fig. 5G,H, Extended Data Fig. 9A,B**), there were significantly higher percentages of BATF-positive CD4^+^ and CD8^+^ T cells (**Fig. 5I,J**) and IFNγ-producing CD4^+^ and CD8^+^ T cells (**Fig. 5K,L**) post Reg-1 KO B7-H3-CAR T cell infusion. No significant differences in other cytokines (IL-10, IL-6, TNFα, IL-17A) or transcription factors (RORγt, TCF1, Foxp3) were observed (**Extended Data Fig. 10A,B**; data for TCF1 and Foxp3 are not shown). These detailed analyses revealed an altered tumor microenvironment elicited by Reg-1 KO B7-H3-CAR T-cell therapy.

### Reg-1 KO B7-H3-CAR T cells induce proinflammatory transcriptional changes in endogenous immune cells

To unbiasedly profile endogenous immune cells, we performed scRNA-seq analysis of the F331 tumor model on day 21 post Ctrl KO or Reg-1 KO B7-H3-CAR T-cell infusion (**Fig. 6A**). This analysis identified 10 endogenous immune cell populations, as well as endothelial cells and fibroblasts (**Fig. 6B**). While all 12 populations were present post Reg-1 KO or Ctrl KO B7-H3-CAR T cell infusion, their composition differed. Specifically, there were decreased frequencies of macrophages and fibroblasts and increased percentages of CD8^+^ T, NK and B cells post Reg-1 KO B7-H3-CAR T-cell infusion (**Fig. 6C**).

**Figure 6:**
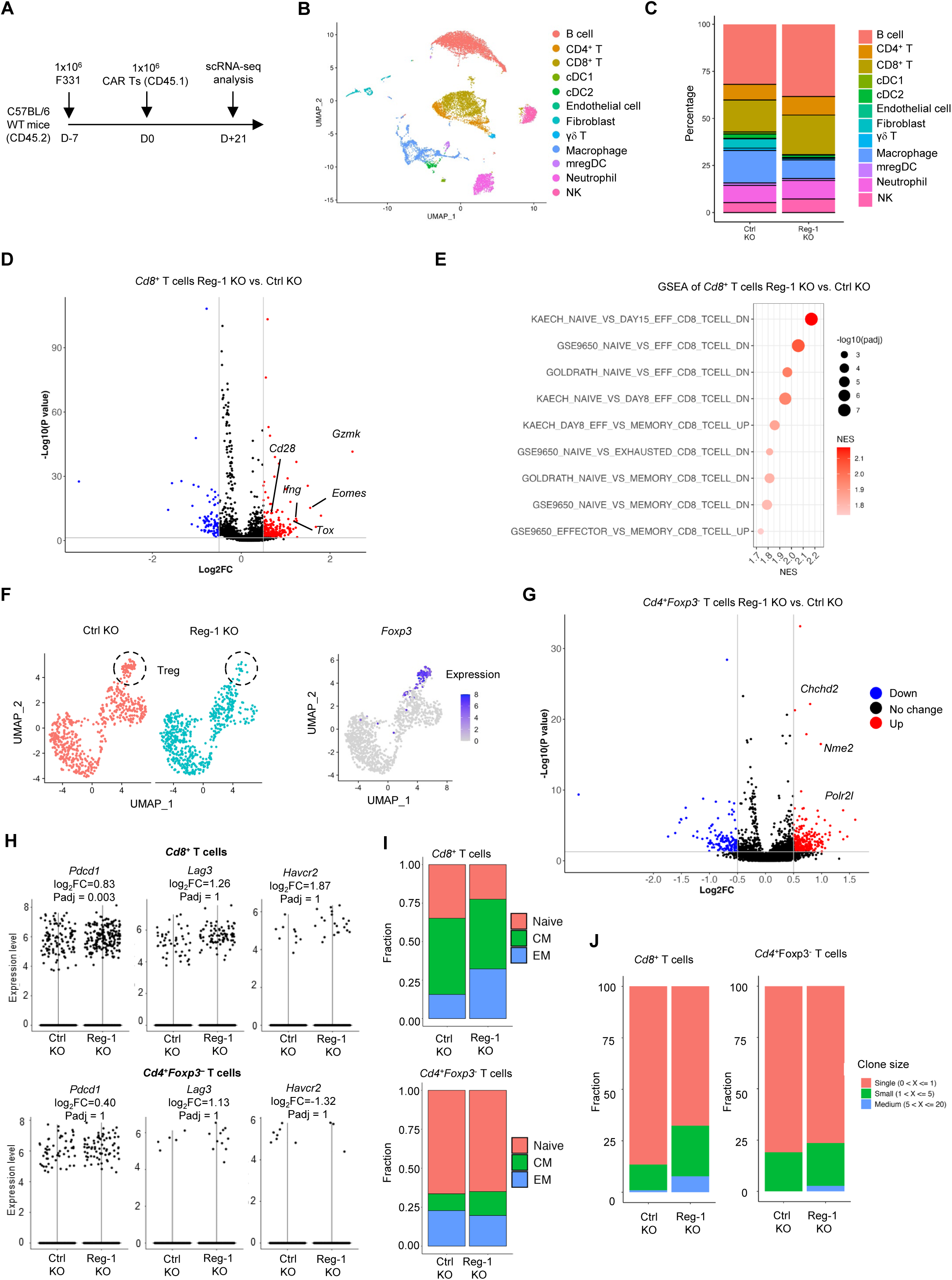
Reg-1 KO B7-H3-CAR T cells increase the effector function on endogenous T and NK cells. (**A**) Experimental schematic: C57BL/6 mice (CD45.2^+^) received a single dose of 1×10^6^ Reg-1 KO or Ctrl KO B7-H3-CAR T cells (Cas9^+^CD45.1^+^) on day 7 post F331 tumor cell injection. Lungs from two mice in each treatment groups were isolated on day 21 post CAR T cell infusion and processed individually for single cell RNA sequencing. (**B**) UMAP indicating clusters of immune cell subsets identified in lungs infiltrating CD45^+^ cells with fibroblast and endothelial cells from Reg-1 KO and Ctrl KO B7-H3-CAR T cells treated mice. (**C**) Percentage of indicated immune cells in lung infiltrating CD45^+^ cells on day 21 post CAR T cell infusion. (**D**) Volcano plot showing upregulated genes in endogenous *Cd8*^+^ T cells in the lungs of Reg-1 versus Ctrl KO B7-H3-CAR T cell treated mice. (**E**) Top 9 significantly (FDR < 0.05) upregulated *Cd8*^+^ T cell related pathways in lung-infiltrating endogenous CD8^+^ T cells from Reg-1 KO versus Ctrl KO B7-H3-CAR T cell-treated mice. (**F**) UMAP projection (left) and *Foxp3* expression (right) in lung-infiltrating endogenous CD4^+^ T cells in Reg-1 KO and Ctrl KO B7-H3-CAR T cell-treated. (**G**) Volcano plot illustrating upregulated genes in lungs infiltrating *Foxp3* negative *CD4*^+^ T cells from Reg-1 KO vs Ctrl KO B7-H3-CAR T cell treated mice. (**H**) Expression of *Pdcd1* (encoding PD-1), *Lag3* and *Havcr2* (TIM-3) on *Cd8*^+^ (top) and *CD4^+^* (bottom) T cells in the lungs of Reg-1 KO vs Ctrl KO B7-H3-CAR T cell treated. (**I**) Fraction of indicated *Cd8*^+^ (top) and *CD4^+^Foxp3^−^* (bottom) T cell subsets in the lungs of Ctrl KO and Reg-1 KO B7-H3-CAR T cell-treated mice. (**J**) Clonal expansion of lungs-infiltrating endogenous CD8^+^ and CD4^+^Foxp3^−^ T cells in Reg-1 KO and Ctrl KO B7-H3-CAR T cell-treated mice.

Transcriptional profiling of endogenous *Cd8*^+^ T cells via scRNA-seq revealed enhanced effector differentiation induced by Reg-1 KO B7-H3-CAR T-cells, as revealed by increased expression of i) effector molecules (*Gzmk*, *Ifng*), ii) costimulatory receptors (*Cd28*), and iii) transcription factors associated with T cell differentiation and/or exhaustion (*Tox*, *Eomes*) (**Fig. 6D**). GSEA revealed enrichments of gene signatures associated with T cell effector differentiation (**Fig. 6E**). UMAP analysis of *Cd4*^+^ T cells revealed that Reg-1 KO B7-H3-CAR T cell-treated mice had a lower frequency of endogenous *Foxp3*^+^ T cells (**Fig. 6F**), while *Cd4*^+^*Foxp3*^−^ T cells highly upregulated key genes in metabolic (*Chchd2*), transcriptional (*Nme2*) and mRNA synthesis (*Polr2l*) pathways (**Fig. 6G**). While endogenous *Cd8*^+^ and *Cd4*^+^ T cells post Reg-1 KO B7-H3-CAR T-cell infusion upregulated *Pdcd1* (PD-1) expression, they did not show altered expression of *Lag3* or *Havcr2* (**Fig. 6H**). Furthermore, Reg-1 KO B7-H3-CAR T cells increased the frequency of endogenous effector *Cd8*^+^ T cells and reduced percentages of naive and memory CD8^+^ T cell populations without alterations in *Cd4*^+^*Foxp3*^−^ T-cell subsets (**Fig. 6I**). Finally, we observed increased clonal expansion of endogenous *Cd4*^+^*Foxp3*^−^ and *Cd8*^+^ T cells post Reg-1 KO B7-H3-CAR T-cell therapy (**Fig. 6J**). These results revealed altered differentiation states of endogenous T cells, including increased effector function in endogenous CD8^+^ T cells, upon Reg-1 KO B7-H3-CAR T-cell therapy.

### Reg-1 KO B7-H3-CAR T cells suppress tumor-promoting signature genes of macrophages

Since our scRNA-seq analysis revealed that Reg-1 KO B7-H3-CAR T cells reduced macrophages and fibroblasts, we next interrogated whether they also induced qualitative changes. Subclustering analysis identified four distinct clusters of fibroblasts (0–3) (**Fig. S14A**), all of which were decreased following Reg-1 KO CAR T-cell therapy (**Fig. 7A,B**). GSEA (**Fig. 7C**) and IPA (**Fig. S14B**) also revealed downregulation of multiple signaling pathways involved in metabolism (e.g., mTORC1) and cytokine (e.g., STAT3) signaling but upregulation of the IFNα response pathway in fibroblasts.

**Figure 7:**
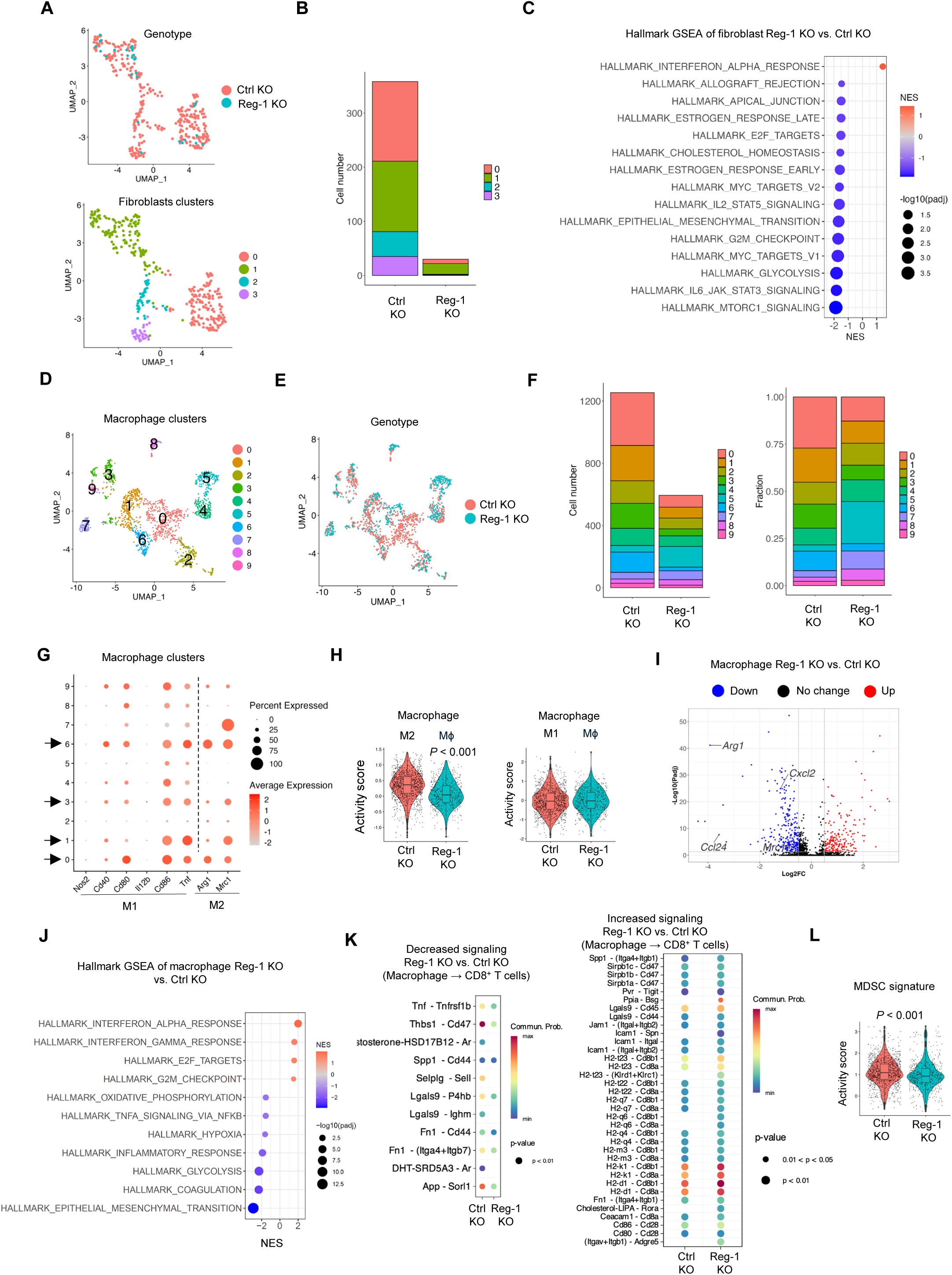
Reg-1 KO B7-H3-CAR T cells suppress tumor-promoting macrophages. (**A**) UMAP projection of fibroblasts in the lungs after infusion with Reg-1 KO or Ctrl KO B7-H3-CAR T cells (n=2); genotype (top) or clusters (botton) are illustrated. (**B**) Absolue cell numbers of fibroblast clusters. (**C**) Significantly (FDR < 0.05) upregulated and top 14 downregulated pathways in fibroblasts, based on GSEA with Hallmark gene sets. (**D**) UMAP projection of 10 unique clusters of macrophages in the lungs of mice on day 21 post Reg-1 KO or Ctrl KO B7-H3-CAR T cell infusion (n=2). (**E**) UMAP projection showing the distribution of macrophages in the lungs of mice post Reg-1 KO or Ctrl KO B7-H3-CAR T cell infusion (color coded by genotype). (**F**) Cell number (left) and fraction (right) of 10 distinct macrophage clusters in the lungs on day 21 post Reg-1 KO or Ctrl KO B7-H3-CAR T cell infusion. (**G**) Dot plot indicating expression levels of M1 or M2 macrophage-associated genes in the clusters. (**H**) Activity scores for M2 (left) and M1 (right) macrophage-related gene signatures in total macrophages from the lungs post Reg-1 KO or Ctrl KO B7-H3-CAR T cell infusion. (**I**) Upregulated and downregulated genes (Log2FC > 0.5 and adjusted p < 0.05) in total macrophages. (**J**) Top 4 significantly (FDR < 0.05) upregulated and top 8 downregulated pathways in total macrophages post Reg-1 KO vs. Ctrl KO B7-H3-CAR T cell infusion, based on GSEA using Hallmark gene sets. (**K**) Decreased and increased signaling pathways identified from the interaction of macrophages and CD8^+^ T cells by CellChat ligand-receptor interaction prediction. (**L**) MDSC signature in macrophages from the lungs showing a significant difference post Reg-1 KO or Ctrl KO B7-H3-CAR T cell infusion.

We next performed subclustering analysis of macrophages and identified 10 distinct clusters (0– 9) (**Fig. 7D**). Reg-1 KO B7-H3-CAR T cells induced a reduction of four (0, 1, 3, 6) clusters (**Fig. 7E,F**). Interrogating all clusters for M1 (*Nos2*, *Cd40*, *Cd80, Il12b*, *Cd86*,*Tnf*)^46^ and M2 (*Arg1*, *Mrc1*)^47^ macrophage marker gene expression revealed that two of the four decreased clusters (clusters 0, 6) expressed *Arg1* (**Fig. 7G**). Among all macrophages, there was also a significant reduction in the activity of an M2 macrophage gene signature (**Fig. 7H**). Furthermore, expressions of *Arg1*, *Mrc1* as well as monocyte-recruiting chemokines (*Ccl24*, *Cxcl2*), was reduced in macrophages from Reg-1 KO B7-H3-CAR T cell-treated mice (**Fig. 7I**). GSEA also revealed upregulated Hallmark IFNα/γ response and cell proliferation pathways (**Fig. 7J**), and CellChat analysis^48^ revealed decreased immunosuppressive Thbs1-CD47 interaction and increased probability of MHCI/II pathway interactions between macrophages and CD8^+^ T cells (**Fig. 7K**). Likewise, gene expression signature analysis revealed a decrease in myeloid-derived suppressor cell (MDSC) signature activity score^49^ in macrophages (**Fig. 7L**). These results collectively indicate that Reg-1 KO enabled B7-H3-CAR T cells to reduce the percentages or function of multiple types of endogenous immunosuppressive myeloid cells.

## DISCUSSION

Here we demonstrate that Reg-1 KO improves the antitumor activity of B7-H3-CAR T cells in immune-competent OS models. Mechanistically, targeting Reg-1 not only improved their effector function but also endowed them with the ability to create a proinflammatory environment characterized by an influx of IFNγ-producing endogenous CD4^+^ and CD8^+^ T cells, as well as NK cells, accompanied by a reduction of inhibitory myeloid cells, including M2 macrophages.

Deleting Reg-1 in B7-H3-CAR T cells enhanced their expansion within 7 days post infusion in lungs and spleens, followed by their contraction within 3 to 4 weeks post infusion. This was associated with transient lymphocytopenia and splenomegaly in both OS models, and monocytopenia in one of the models. While CAR T-cell expansion and splenomegaly were dependent on B7-H3-CAR expression, transient lympho- and monocytopenia also occurred post Reg-1 KO T cells that expressed a CAR specific for an irrelevant antigen (SP6). Cytokine and chemokine measurements post generation of Reg-1 KO SP6- and B7-H3-CAR T cells did not show a unique signature. Likewise, there was no common cytokine/chemokine signature in the peripheral blood of mice on day 7 post infusion. However, mice that received Reg-1 KO B7-H3-CAR T cells had significantly higher levels of chemokines (MCP-1, MIG, MIP-2) and cytokines (IL-6, IFNγ). Of these chemokines and cytokines, only IFNγ was expressed at significantly higher levels by Reg-1 KO B7-H3-CAR T cells compared to Ctrl-KO B7-H3-CAR T cells, highlighting the contribution of endogenous immune cells to creating an inflammatory environment post CAR T-cell therapy. IL-6 and IFNγ are hallmark cytokines of immune effector cell-associated side effects such as cytokine release syndrome (CRS)^50–54^; however, none of the mice post Reg-1 KO B7-H3-CAR T cells developed any clinical signs of distress or lost weight.

Transcriptional profiling of Reg-1 KO B7-H3-CAR T cells on day 5 post infusion was consistent with our previous finding for CD19-CAR and TCR-transgenic T cells, including upregulation of transcription factors such as BATF and proinflammatory and metabolic signaling pathways^10,11^. In addition, we found increased expression of *Ifng* and *Pou2f2*, consistent with recent findings in Reg-1 KO NK cells^23^. Finally, Reg-1 KO induced a recently described CXCR6-positive CD8^+^ T cell expression signature that counterbalances PD1-mediated immune suppression^44,55^. Flow cytometric analysis revealed increased EM differentiation of Reg-1 KO versus Ctrl KO B7-H3-CAR T cells. Post expansion, all CAR T cells, regardless of Reg-1 expression, contracted to low levels by day 21, and a significant number of these CAR T cells expressed PD-1 and PD-L1, suggestive of signaling in cis as described for antigen presenting cells^56^. Despite being able to detect CAR T cells by flow cytometry and IHC, we could not detect CAR T cells by scRNA-seq at this late time point, precluding a detailed transcriptomic analysis. The observed contraction of B7-H3-CAR T cells was greater than we previously reported for CD19-CAR T cells in immune-competent leukemia animal models^11^, suggesting that persistence of Reg-1 KO CAR T cells is influenced by the tumor type and the associated microenvironment they are targeting.

Our endogenous immune cell analysis of PBS-treated tumors was consistent with the immune cell profile of OS lung metastasis, demonstrating a high percentage of myeloid cells, including M2 macrophages^57^. On day 7 post infusion, Reg-1 KO B7-H3-CAR T cells had significant effects on endogenous immune cells in both OS models, including an influx of CD8^+^ T cells and NK cells into tumor-bearing lungs. While changes in the percentage of myeloid cells in lungs and spleens were model dependent, there was a consistent increase of PD-L1 expression on M2 macrophages and DCs, raising the possibility that endogenous myeloid cells may potentially dampen the inflammatory response induced by Reg-1 KO CAR T cells. However, the flow cytometric analysis on day 21 revealed a decrease in M2 macrophages, coinciding with an increase in M1 macrophages. This indicates a Reg-1 KO CAR T cell-induced reversal of the immunosuppressive environment observed on day 7, which might explain why additional PD-L1 blockade did not further improve their antitumor activity in our model. Limited benefit of combining checkpoint blockade with CAR T cells has also been reported in preclinical and early phase clinical studies^58,59^. Our transcriptomic profiling of endogenous immune cells on day 21 after Reg-1 KO CAR T-cell infusion confirmed and extended upon our flow cytometric analysis. For example, pathway analyses revealed enrichments of gene signatures associated with effector differentiation in T cells upon Reg-1 KO CAR T-cell therapy, while macrophages showed a significant reduction in M2 macrophage and MDSC gene signatures. One notable aspect of our flow cytometric and transcriptomic analyses was that these effects were present at a time when Reg-1 KO CAR T cells had contracted. In this regard, future studies are needed to delineate in greater detail the temporal changes in the endogenous immune landscape following Reg-1 KO CAR T-cell therapy.

While Reg-1 KO in CAR T cells improved their antitumor activity, tumors eventually progressed. We excluded the development of antigen-loss variants as a potential mechanism of failure, and so tumor progression is most likely linked to the contraction of CAR T cells within 3 weeks post infusion. One explanation might be that we did not use lymphodepleting chemotherapy like cyclophosphamide in our study. While it is routinely used prior to CAR T-cell infusion^60–63^, it was not used here, because it has direct antitumor effects and also transiently depletes endogenous immune cells. Other investigators have demonstrated that deleting Roquin-1 or BCOR further enhances the effector function of Reg-1 KO CAR T cells^24,25^. The benefit of combining these or other genetic modifications remains to be tested in our OS models for Reg-1 KO B7-H3-CAR T cells in the future.

In conclusion, we demonstrate that deleting the negative regulator Reg-1 KO improved the effector function of B7-H3-CAR T cells, and more importantly, endowed them with the ability to create a proinflammatory environment. Others have reported that expression of secretory cytokines can remodel the tumor microenvironment^64,65^, and likewise, CAR design-dependent remodeling of the TME has been reported^66^. However, to our knowledge, this is one of the first reports demonstrating such effects for a negative regulator of T cell function. Thus, our findings highlight that deleting negative regulators in CAR T cells presents a promising approach to target the inhibitory immune landscape of solid tumors.

## ACKNOWLEDGMENTS

We would like to acknowledge Nicole Chapman and Jana Raynor for reviewing the manuscript; Amanda George for assistance with the *in vivo* mouse studies; Melissa Hendren for Cas9 mice colony management and the staff of the Center for Advanced Genome Editing (CAGE) for performing indel analyses. We appreciate the help of the Department of Immunology Flow Cytometry core for cell sorting of samples used for scRNA-sequencing. We would also like to thank the staff of the Hartwell Center for Biotechnology for sequencing, and the Comparative Pathology Core for IHC and ISH analyses, which are both in part supported by National Institute of Health (NIH)/National Cancer Institute (NCI) grant P30CA021765. This work was supported by NIH/NCI grant 1T32CA272387 to A.O.A., R35CA253188, U01CA281868 and R01AI140761 to H.C., and R01CA292466 and 1R01CA288967 to J.T.Y. Additional support was provided by the American Lebanese Syrian Associated Charities (ALSAC) to G.K., C.D., H.C., S.G. The content is solely the responsibility of the authors and does not necessarily represent the official views of the NIH.

## Author Contributions

Conceptualization: A.O.A., H.C., S.G.; Methodology: A.O.A., H.S., S.S.P., T.C., H.S, P.Z., S.M.P.; Investigation: A.O.A., H.S., P.N., S.S.P., T.C., X.S., P.Z., H.S., A.K., M.S.P.; Resources: J.T.Y., H.C., S.G.; Formal analysis: A.O.A., H.S., S.S.P., H.S., J.Y.M., L.T., S.G.; Supervision: H.C., S.G.; Funding acquisition: G.K., C.D., H.C., S.G.; Writing – original draft preparation: A.O.A., S.G.; Writing – review and editing: A.O.A., H.S., S.S.P., H.S., P.N., X.S., P.Z., J.Y.M., T.C., A.K., L.T., V.P., M.S.P., D.M.L., S.M.P., J.T.Y., G.K., C.D., H.C., S.G.

## METHODS

### Cell lines

The murine F331 (Col2.3-Cre, p53^fl/+^) and M1199 (Col2.3-Cre; TP53fl/fl; LSL-cMycT58A) OS cell lines were derived from genetically engineered mouse models (GEMM) that previously have been published^67^. Murine tumor cell lines were cultured in DMEM media (GE Healthcare Life Sciences, Chicago, IL) supplemented with 10% fetal bovine serum (FBS), 1% glutamax and 1% penicillin-streptomycin. ATCC short-tandem repeat profiling service was used to authenticate cell lines. Cells lines were routinely screened for mycoplasma by using MycoAlert Mycoplasma Detection kit (Lonza, Walkersville, MD). Cell lines were maintained in culture for a maximum of 2 months before being replaced with a fresh vial of cells. B7-H3 KO F331 cells (F331-KO) were generated using CRISPR/Cas9 technology at St. Jude Center for Advanced Genome Engineering (CAGE) as previously described using an established protocol^27,68^ with precomplexed ribonuclear proteins (RNPs) using a sgRNA targeting B7-H3 (CAGE193.CD276.g22, 5’-UGUCCACCAGGGCCACCACG 3’, Synthego). Cells were clonally expanded and screened for the desired modification (out-of-frame indels and a premature stop codon) via targeted deep sequencing using gene specific primers with partial Illumina adapter overhangs (CAGE193.DS.F – 5’-CTACACGACGCTCTTCCGATCTacctgaaaggaagcgtgtgcagagc-3’ and CAGE193.DS.R-5’-CAGACGTGTGCTCTTCCGATCTagctgccctcgtcggttactcggac-3’, overhangs shown in uppercase) as previously described^68^. NGS analysis of clones was performed using CRIS.py^69^. Final clones were authenticated using the PowerPlex® Fusion System (Promega) performed at St. Jude Hartwell Center for Biotechnology. Knockout was confirmed by flow cytometry analysis.

### Animal models

All animal experiments were performed according to an approved protocol by St. Jude Children’s Research Hospital Institutional Animal Care and Use Committee. Eight-ten-week-old male and female mice maintained under ambient temperature and humidity with a 12/12 hour (h) on/off light cycle were used for the F331 and M1199 models. C57BL/6 and Rosa26-Cas9 knock-in^70^ mice were purchased from Jackson Laboratory. For the F331 model, 1×10^6^ cells were injected intravenously (i.v.) on day 0 into C57BL/6 mice. 1×10^6^ of either Reg-1 KO SP6-, Ctrl KO B7-H3-, Reg-1 KO B7-H3-CAR T cells in sterile PBS were injected i.v. on day 7. In the M1199 model, 2×10^5^ cells were injected i.v. on day 0 into C57BL/6 mice. 1×10^6^ of either Reg-1 KO SP6-, Ctrl KO B7-H3-, Reg-1 KO B7-H3-CAR T cells in sterile PBS were injected i.v. on day 7. Weekly body weight measurements and routine health checks were performed. Subsets of mice were bled retro-orbitally to collect blood for complete blood count analysis (Genesis ^TM^ analyzer) and cytokine/chemokine analysis (Luminex Corporation). The study endpoint was attained either when mice lost about >20% of initial body weight, or showed signs of disease or discomfort. To determine antitumor activity, mice were euthanized on day 21 (F331) or day 28 (M1199) post T cell infusion.

For subcutaneous F331 model, mice received 2×10^6^ F331 cells in the right flank. On day 7, mice were administered a single i.v dose of either 5×10^6^ Ctrl KO- or Reg-1 KO B7-H3-CAR T cells via tail vein injection. Tumor kinetics were monitored by weekly caliper measurements. Study endpoint was attained when mice i) lost 20% of body weight, ii) showed signs of distress iii) tumor burden reached 20% of total body mass (≥ 2,500 mm^3^) or iv) when recommended by veterinary staff.

For flow cytometric and scRNA seq analyses, mice were euthanized and single cell suspension of lungs and spleens were prepared. Briefly, lungs and spleens were isolated from osteosarcoma bearing mice and single cell suspensions of lungs were prepared by mechanical dissociation and enzymatic digestion with collagenase IV (0.5 mg/mL, STEMCELL Technologies) and DNase I (25 IU/mL, Sigma Aldrich). Cell suspension was incubated at 37°C for 30 mins and then passed through a 70 µm filter to remove debris. Cell supernatants were spun to obtain cell pellets. Ammonium chloride (ACK) lysing buffer (Gibco) was added to the cell pellet to remove red blood cells. Spleens were only mechanically dissociated and passed through a 70µm filter before red blood cell lysis. Cells were washed with PBS and resuspended in flow cytometry staining buffer.

#### Generation of retroviral vectors

The generation of the retroviral vectors encoding B7-H3- or SP6-CAR.CD28.z was previously described^71^.^72^ To generate retroviral particles, GPE+86 producer cell line was cultured in IMDM media supplemented with 10% FBS and 1% glutamax to generate retroviral particles for murine T cell transduction. Cell supernatant was collected after 48 hours, spun at 1,000g for 5 mins and passed through a 0.45μm filter. Fresh supernatants or leftover supernatant snap-frozen at -80 °C was used for chimeric antigen receptor (CAR) T cell transduction. The generation of the retroviral vectors encoding sgRNAs (non-targeting control (Ctrl) sgRNA, ATGACACTTACGGTACTCGT or Reg-1 sgRNA, AAGGCAGTGGTTTCTTACGA) and ametrine were previously reported^10^. To generate retroviral particles, the Platinum E retroviral packaging cell line (Cell Biolabs, Inc.) was co-transfected with sgRNA-encoding retroviral vectors and helper plasmid pCL-Eco (Addgene no. 12371).

#### Generation of murine CAR T cells

Naive CD8^+^ T cells were enriched from the spleen and lymph nodes of Cas9 (CD45.1) transgenic C57BL/6 mice using a naive CD8 T cell isolation kit (Miltenyi Biotec, Bergisch Gladbach, Germany). Enriched naive CD8^+^ T cells were activated in the presence of murine CD3 (1μg/mL, BD Biosciences) and CD28 (2μg/mL, BD Biosciences) antibodies and cultured with human IL2 (50IU/mL, PeproTech) in complete RPMI media for 48h. Reg-1 or non-targeting single guide RNA (Ctrl) KO B7-H3-CAR or Reg-1 KO SP6-CAR T cells were generated by co-transduction of activated naïve CD8^+^ T cells with retroviral vectors encoding CARs or the respective guide RNA + ametrine. Cells were cultured in the presence of human IL2 (50IU/mL, PeproTech) for 48h while non-transduced (NT) naive CD8^+^ T cells served as a control. Cells were expanded in the presence of IL7 (10ng/mL, PeproTech) and IL15 (10ng/mL, Peprotech) for another 2-3 days. Editing efficacy was confirmed by indel analysis and double transduction by flow cytometry. CAR T cells were used for experiments on or before day 6 post-transduction.

#### Multiplex analysis

Cell culture supernatants from murine CAR T cells and non-transduced T cells were collected 4 days post-transduction or obtained from a co-culture of CAR T cells and tumor cells at an effector to target ratio of 1:1 after 48 h. Serum was obtained from tumor-bearing mice that received CAR T cell therapy or PBS on day 7. All samples were snap-frozen at -20°C before evaluating cytokine/chemokine production with a 32-plex Milliplex mouse cytokine/chemokine magnetic bead panel (MCYTMAG-70K-PX32, Millipore Sigma). Analysis was performed using a Luminex FlexMap 3D instrument and software (Luminex Corporation).

#### Cytotoxicity assays

CAR T cell cytotoxicity was assessed with a CellTiter96 AQueous One Solution Cell Proliferation Assay kit (Promega). In a 96-well tissue culture treated plate, 10,000 F331 cells and 8,000 M1199 cells were co-cultured with serial dilutions of CAR T cells and incubated at 37°C for 48 h. Media, CAR T cells and tumor cells alone served as controls. Each condition was performed in technical triplicates. The media and CAR T cells were removed by slowly pipetting up and down without disturbing the adherent tumor cells. CellTiter96 AQueous One Solution Cell Proliferation reagent in complete DMEM was added into each well in the plate and incubated at 37°C for 2 h. The percentage of viable tumor cells after exposure to CAR T cells was calculated by the formula: (Sample-media alone)/(Tumor alone -media alone) x 100.

#### Flow cytometry

Cell surface staining was conducted using 5% FBS in PBS (Lonza) as the antibody diluent and cell wash medium. Cells were labeled at 2×10^6^ cells per mL with optimally titrated antibody solutions and incubated at 4°C for 30 min. Unless noted otherwise, dead cells were excluded through reaction with Live/Dead Fixable Blue dead cell stain (Thermofisher) according to the manufacturer’s protocol, except where otherwise stated.

#### B7-H3 and CAR detection

Matched isotypes or known negatives (such as non-transduced (NT) T cells) served as gating controls. B7-H3 detection was performed using anti-mouse CD276-APC (1:200 dilution, clone MIH32, BD Biosciences). CAR expression was determined using anti-mouse F(ab’)2 fragment specific antibody (1:200 dilution; polyclonal, Jackson ImmunoResearch). Dead cells were excluded by staining cells with 4’,6-diamidino-2-phenylindole (DAPI) before data acquisition. A FACSCanto II instrument (BD Biosciences) equipped with 405, 488, and 633nm lasers, and running BD FACS DIVA v.9.0 software was used to acquire the flow cytometry data.

#### Cell surface staining of single cell suspensions from lungs and spleens

Cells were stained with fluorochrome-conjugated antibodies using combinations of the markers shown in Tables S1 and S3. Flow cytometric analysis was conducted using a 5-laser Cytek Aurora cytometer (Cytek Biosciences, Fremont, CA, USA) equipped with 355, 405, 488, 561, and 633 nm lasers and 56 fluorescent detectors, utilizing SpectroFlo software (Cytek) for data acquisition and spectral unmixing. Dimensional reduction and additional analyses were conducted on unmixed FCS files using the OMIQ online software suite (Dotmatics, Boston, MA, USA).

#### Intracellular cytokine staining of single cell suspensions from lungs

Cells were stimulated with ionomycin calcium salt (1 μg/mL, Sigma-Aldrich) and phorbol 12-myristate 13-acetate (PMA, 50 ng/ml, Sigma-Aldrich) for 1 hour. Brefeldin A (2μL of 1000×) was then added for an additional 3 hours to block cytokine secretion. Stimulation and blocking of protein transport were performed at 37°C in 5% CO2. Cells were collected from culture and washed in staining buffer at 4°C, then labeled for surface epitope expression as described above. Afterward, cells were fixed and permeabilized using the BD Cytofix/Cytoperm kit (BDBiosciences) as per product instructions. Immunofluorescent staining against intracellular antigens was performed. Surface and intracellular staining was conducted with antibodies detailed in Table S5. Flow cytometric data was acquired using a Cytek Aurora cytometer described above and spectrally unmixed FCS files were analyzed using the Flow jo software (v10.8.1),

#### scRNA-seq sample preparation and data analysis

#### ScRNA-Seq sample preparation and bioinformatics analysis

Ctrl or Reg-1 KO CAR T cells were infused into F331-tumor bearing mice (see animal models section). Lung and spleen were isolated and processed for single cell suspension. In the experiment to profile CAR T cells on day 5 post infusion, lungs or spleens from three mice per treatment group were pooled together. Ctrl or Reg-1 KO CAR T cells were sorted based on CD45.1, CD8 and Ametrine expression. To investigate the lung OS immune landscape 21 days post CAR T cell infusion, two mice per treatment group were euthanized and lung was processed individually. Sorted CD45-positive and CD45-negative cells from lung Ctrl or Reg-1 KO at a ratio of 3:1 was processed for scRNA-seq experiment. Dead cells were excluded by staining with DAPI before acquisition. Cells were sorted on a BD FACS Aria III cell sorter (BD Biosciences). The samples were centrifuged at 2,000 rpm for 5 minutes, after which cells with over 98% viability were resuspended in 1× PBS (Thermo Fisher Scientific) containing 0.04% BSA (Amresco), at a final concentration of 1 × 10^6^ cells per ml. Single-cell suspensions were then loaded into a Chromium Controller (10× Genomics) to produce approximately 10,000 single-cell gel bead emulsions per sample, with each sample processed in a separate channel. Library preparation was carried out using the Chromium Next GEM Single Cell 5′ v2 (dual index) and Gel Bead kit (10× Genomics). Amplified cDNA (11 cycles) was either used for TCR enrichment and library construction with the Chromium Single Cell V(D)J TCR kit or for gene expression library preparation. The quality of the cDNA and resulting libraries was assessed using a high-sensitivity DNA chip on a 2100 Bioanalyzer (Agilent Technologies). Sequencing of the 5′ libraries was performed on an Illumina NovaSeq system, with paired-end reads consisting of 26 cycles for read 1 and 90 cycles for read 2. On average, gene expression libraries yielded 50 million reads per sample, while TCR libraries generated 5 million reads per sample.

#### Data preprocessing

The Cell Ranger v6.0 Single-Cell software suite (10× Genomics) was used to process the scRNA-seq FASTQ files. The ‘cellranger count’ command was performed to align the raw FASTQ files to a custom mouse reference genome (mm10 with the addition of the mouse B7-H3-CAR sequence) and summarize the data into matrices that describe gene read counts per cell. For the datasets with matched TCR sequencing data, the ‘cellranger vdj’ command was used to generate a count matrix, which, after filtering, was used for downstream analyses.

For gene expression sequencing, the filtered count matrices were read into the R package Seurat (v4.1). Samples were merged into single Seurat objects for consistent filtering, and features detected in fewer than three cells were removed from the dataset. Cells with abnormally low features or unique molecular identifier (UMI) counts or high mitochondrial read percentages (potentially dead or damaged cells) were removed. Cells with abnormally high UMI counts (potentially multiple cells in a single droplet) were also removed. Finally, any remaining multiplets expressing mutually exclusive marker genes were removed.

For Ctrl or Reg-1 KO CAR T cells from lungs and spleens on day 5 post infusion, 7,282 lung derived CAR T cells were retained with an average of 3,533 genes per cell (UMI median: 13,228; range: 1,581–59,735) and 5,059 spleen derived CAR T cells were retained with an average of 3,444 genes per cell (UMI median: 11,336; range: 1,477–59,877).

For tumor microenvironment cells on 21 days post Ctrl or Reg-1 KO CAR T cell infusion, 13,255 cells were retained with an average of 1,933 genes per cell (UMI median: 4,495; range: 500– 49,880). After quality control, libraries were normalized with the Normalize Data function (scale factor = 1×10^6^) in the Seurat R package.

For TCR sequencing, filtered contig annotation matrices from the Cell Ranger output were loaded into R. Annotation and quantification of TCR clonotypes were processed with the scRepertoire package (v1.3.5). Clonal frequencies were categorized as follows: single (1), small (2–5) and medium (6–20).

#### Cluster annotation and data visualization

Normalized and filtered data were processed using the standard Seurat pipeline (v4.1). UMAP dimensionality reduction was used for visualization, and Seurat’s FindClusters function (v4.1) was used to separate cells into unsupervised clusters. Cell types in clusters were defined using the following marker genes: CD8^+^ T cells (*Cd3d*^+^*Cd8a*^+^), CD4^+^ T cells (*Cd3d*^+^*Cd4*^+^), γδ T cells (*Cd3d*^+^*Tcrg-V6*^+^), B cells (*Cd19*^+^ *Cd79a*^+)^, NK cells (*Ncr1*^+^), macrophages (*Adgre1*^+^), cDC1 (*Xcr1*^+^), cDC2 (*Cd209a*^+^), mregDC (*Ccr7*^+^*Cd200*^+^), neutrophils (*Hdc*^+^), Fibroblasts (*Pdgfra*^+^), Endothelial cells (*Cdh5*^+^). For analysis of CD8+ T cells, *Cd3d*^+^*Cd8a*^+^*Cd4*^-^ cells were isolated and re-clustered using the Seurat workflow.

#### Identification of ligand–receptor pairs in scRNA-seq data

To determine the differences in cell-cell interaction strengths in tumor microenvironment between Ctrl or Reg-1 KO CAR T cell treated lung, CellChat (v1.4.0) was used. CellChat objects were produced using the CreateCellChat function. The overall cell–cell interaction strengths between different cell types was calculated by the default workflow based on ligand–receptor databases in the CellChat package. After running the pipeline on the two genotypes separately, the two CellChat objects were merged using the mergeCellChat function. Heat maps of the differential cell–cell interaction strength between genotypes were generated using the function netVisual_heatmap (measure = ‘weight’), with row z score to indicate the difference in the interaction strength on the heat map.

#### Differential gene expression analysis

Differential gene expression analysis was performed on the log-normalized gene expression matrices with the Seurat FindMarkers function (v4.1) by using a two-tailed Wilcoxon rank-sum test and calculating adjusted *P* (*P*adj) values via Bonferroni correction.

#### GSEA analysis

For GSEA, genes were ranked in order of descending log (fold change) values (derived from the differential expression analysis). Pre-ranked GSEA was performed using the broad GSEA command line software (v4.0.3). The Hallmark, C2 and C5 geneset collections from MSigDB (v7.3) were used.

#### IPA upstream regulator inference

Top 250 genes for up- and down-regulated genes (500 genes in total) in each differential gene expression comparison are used as the input for Ingenuity Pathway Analysis (IPA, Qiagen) after the Log2 fold change cut-off, followed by “Core Analysis”.

### Pathological examination

Mice injected with F331, M1199 or B7-H3 KO F331 were euthanized at different timepoints post T cell infusion and submitted to the St. Jude Children’s Research Hospital Comparative Pathology Core Laboratory for histological assessment to ascertain the extent of metastatic disease and evaluate pulmonary microenvironment. The mice were necropsied following euthanasia and lungs were immersed in 10% neutral-buffered formalin for at least 48 hours. After fixation, tissues were embedded in paraffin, sectioned at 4 μm, mounted on positively charged glass slides (Superfrost Plus, Thermo Fisher Scientific), and dried at 60°C for 20 minutes before dewaxing and staining with H&E. Pathology evaluation was conducted in a blinded manner to the experimental condition. HE, IHC, and ISH images were scanned with a Pannoramic 250 Flash III Scanner (3DHistec) and imaged and analyzed using HALO v3.6.4134 and HALO AI 3.6.4134 (Indica Labs, Inc. Albuquerque, NM). Antibodies used for IHC are listed in **Table S6**.

### In-situ Hybridization (ISH)

H&E-stained section of the lungs and spleens were immunolabeled with a customized RNAscope™ 2.5 VS probe set to detect murine B7-H3-CARs (CAR-scFv-m276-C1, Cat No.1220899-C1, Advanced Cell Diagnostics Inc., Hayward CA). Staining was performed on a Ventana Discovery Ultra automation system (Ventana Medical Systems, Inc., Tucson, AZ) using the RNAscope VS Universal AP Reagent Kit (RED) (Advanced Cell Diagnostics, Inc., Hayward, CA) and the following reagents from Roche Diagnostics mRNA RED Detection Kit (760-234), mRNA Sample Prep Kit (760-248), and mRNA RED Probe Amplification Kit (760-236) according to manufacturer’s protocols. The hybridization signals were detected by chromogenic development with fast red, followed by counterstaining with hematoxylin. Each sample was quality controlled for RNA integrity with a RNAScope probe specific to murine *Ppib* RNA (313919, Advanced Cell Diagnostics, Inc., Hayward, CA) and for background with a probe specific to bacteria gene *dapB* RNA (312039, Advanced Cell Diagnostics, Inc., Hayward, CA). Specific RNA staining signal was identified as red punctate dots. Pathology evaluation was conducted in a manner that was blinded to the experimental condition.

### Statistical analysis

The sample numbers (n), biological replicates, *P* value and statistical methods are shown in the figure legends. GraphPad Prism software v.10.4.0 was used to plot all graphs and perform statistical analyses. Data are presented as mean ± standard error of mean (SEM).

## Data availability

The authors declare that the data supporting the findings of this study are available within the manuscript and its Supplementary Information. For material requests, contact the corresponding authors. All scRNA-seq data described in the manuscript have been deposited in the NCBI GEO database.

## Extended Data Figures

**Extended data Figure 1:**
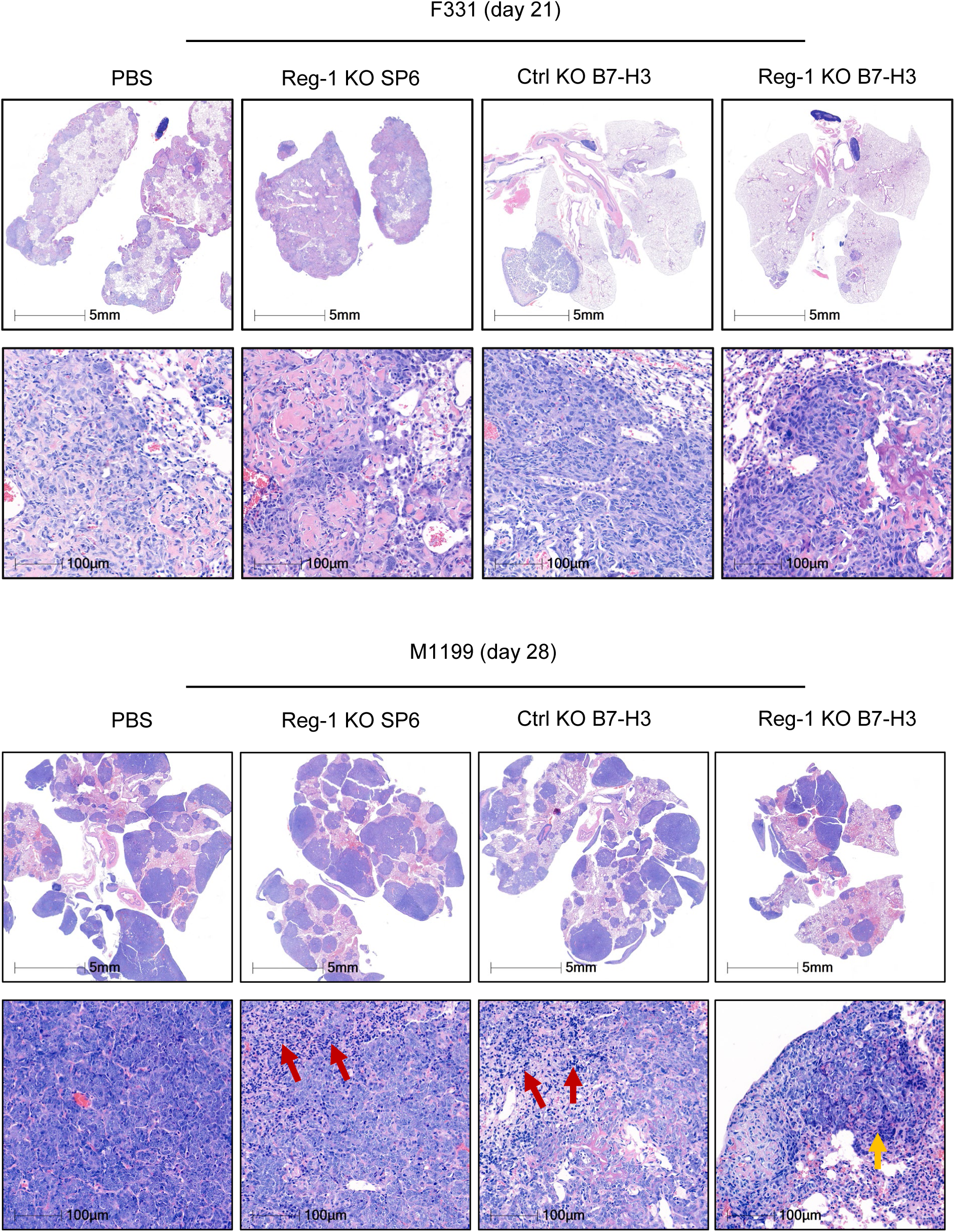
H&E staining of osteosarcoma pulmonary tumor post-CAR T cell infusion. Hematoxylin and eosin (H&E) staining of lungs from mice on day 21 (F331; upper) and day 28 (M1199; lower) after infusion with Reg-1 KO SP6, Ctrl KO or Reg-1 KO B7-H3-CAR T cells or PBS. Representative images. Top rows: 0.6× magnification, 5 mm scale; bottom rows: 20× magnification, 100 µm scale.

**Extended data Figure 2:**
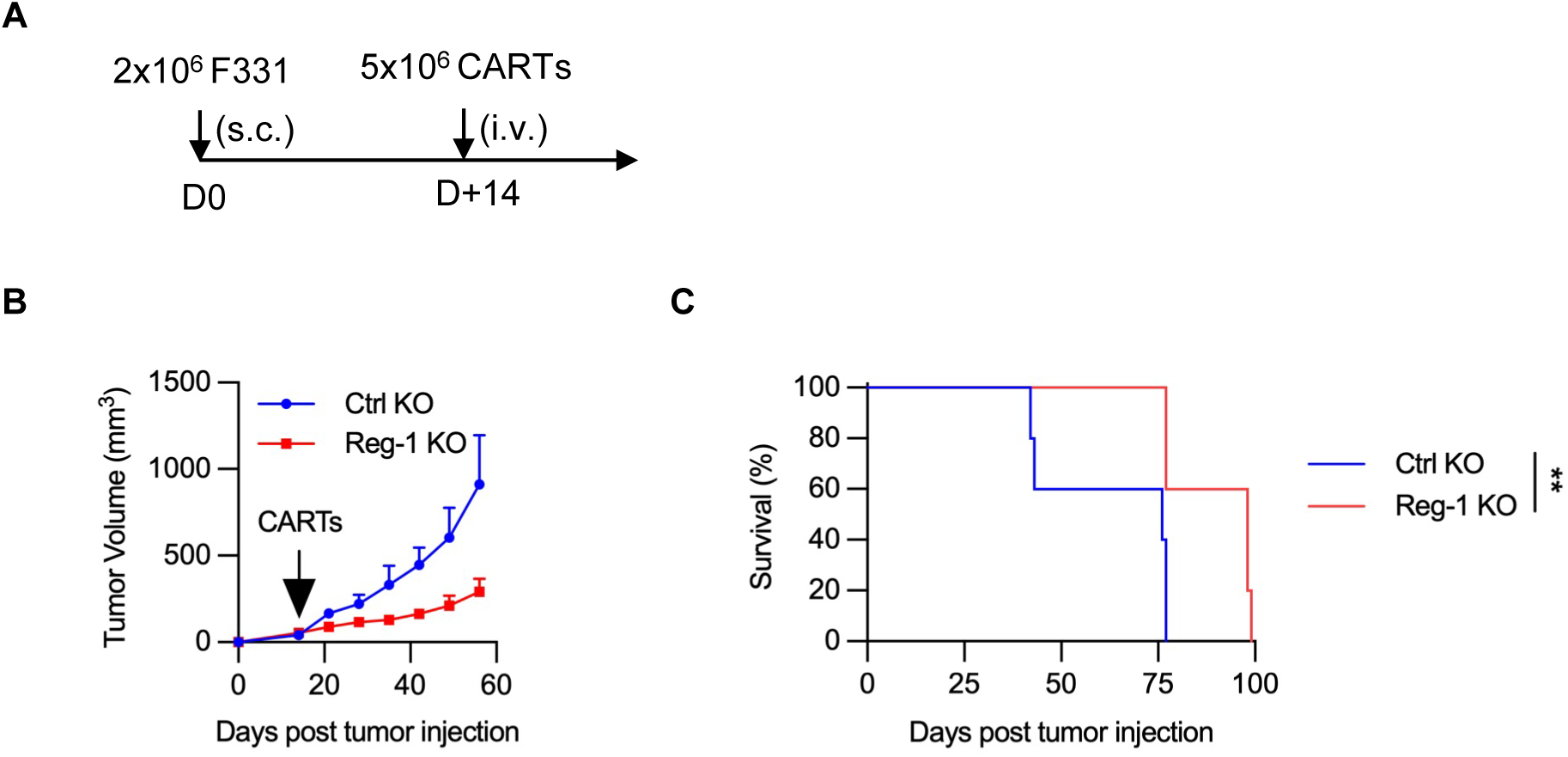
Reg-1 KO improves antitumor activity of B7-H3-CAR T cells in subcutaneous mouse model. (**A**) Experimental schematic: C57BL/6 mice were implanted with 2×10^6^ F331 subcutaneously on the right flank and on day 14 received a single i.v. dose of 5×10^6^ Reg-1 KO or Ctrl KO B7-H3-CAR T cell infusion. (**B**) Tumor growth curve (n=5 mice per group). (**C**) Kaplan-Meier survival curve, log-rank Mantel-Cox test, n=5 mice per group, **:p<0.01.

**Extended data Figure 3:**
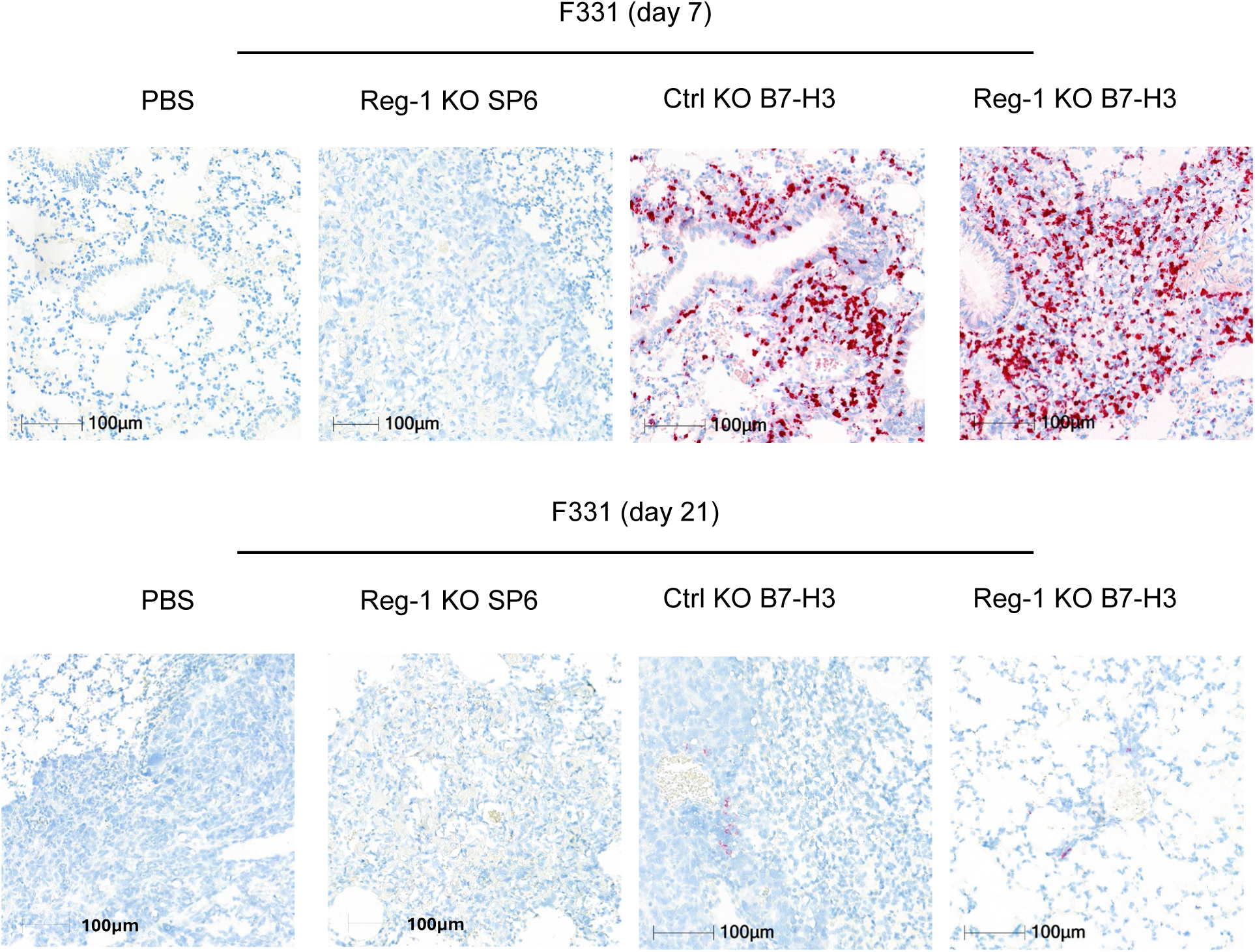
B7-H3-CAR T cells are present within pulmonary metastases on day 7 post-infusion. Detection of B7-H3-CAR T cells with an *in situ* hybridization (ISH) probe specific for the B7-H3-CAR in the F331 model on day 7 (top) and day 21 (bottom) after infusion with Reg-1 KO SP6, Ctrl KO or Reg-1 KO B7-H3-CAR T cells. Fast red chromogen with hematoxylin counterstain. 20× magnification, 100 µm scale.

**Extended data Figure 4:**
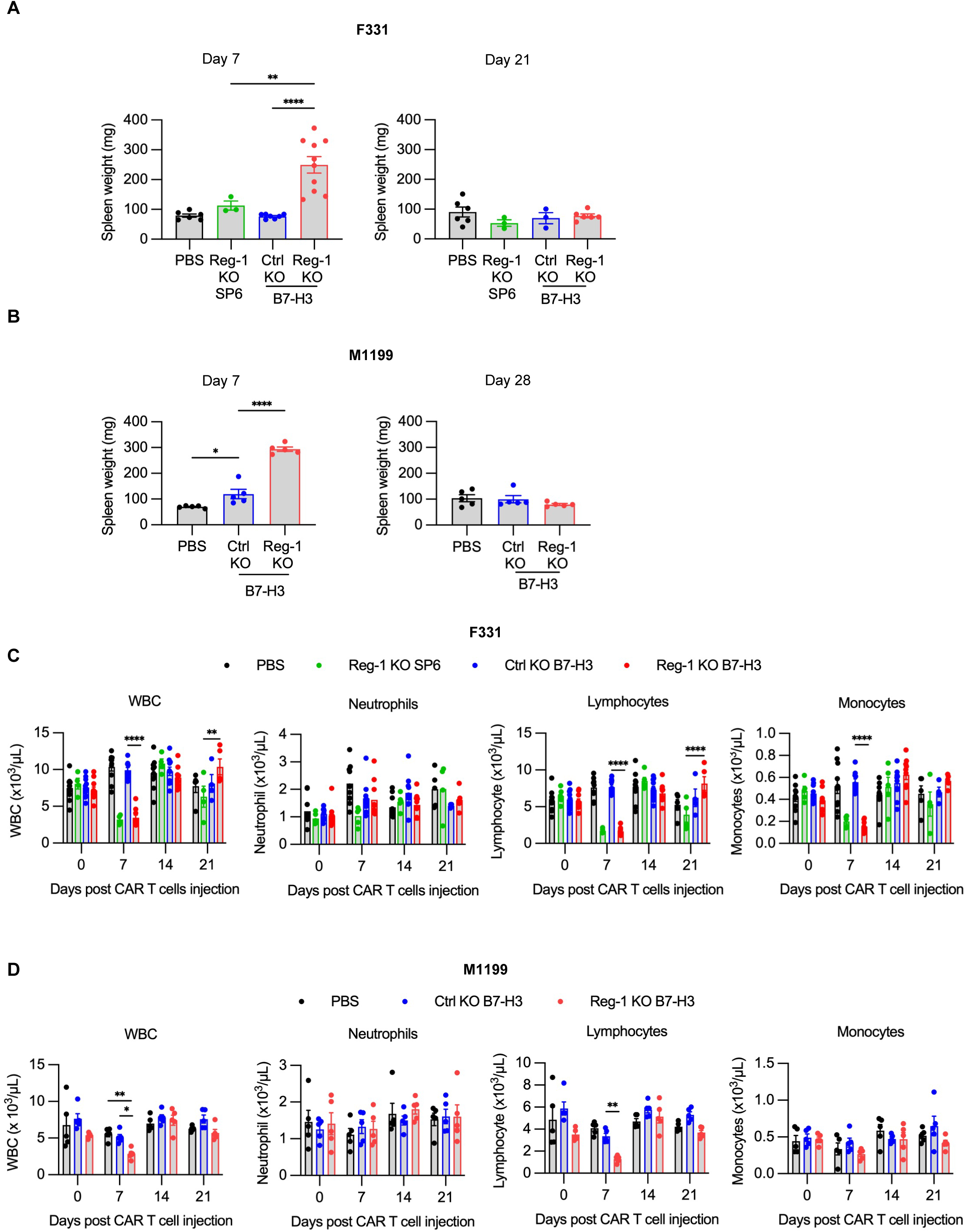
Reg-1 KO B7-H3-CAR T cells induced transient splenomegaly, lympho- and monocytopenia on day 7 post injection. (**A**) Mice were injected with F331 and received indicated CAR T cells as described for **Fig. S9A**. Spleens weight is shown for days 7 and 21 post CAR T cell infusion (n=3-10 mice per group). (**B**) Mice were injected with M1199 cells and received indicated CAR T cells as described for **Fig. S9B**. Weight of spleens on days 7 and 28 post CAR T cell infusion (n=5 mice per group). (**C,D**) Complete blood count analysis on days 0, 7, 14 and 21 post-CAR T cell infusion in the (**C**) F331 or (**D**) M1199 model (n=5-10 mice per group) One way or Two-way ANOVA with Tukey’s test for multiple comparisons; *: p<0.05; **:p<0.01; ****:p<0.0001.

**Extended data Figure 5:**
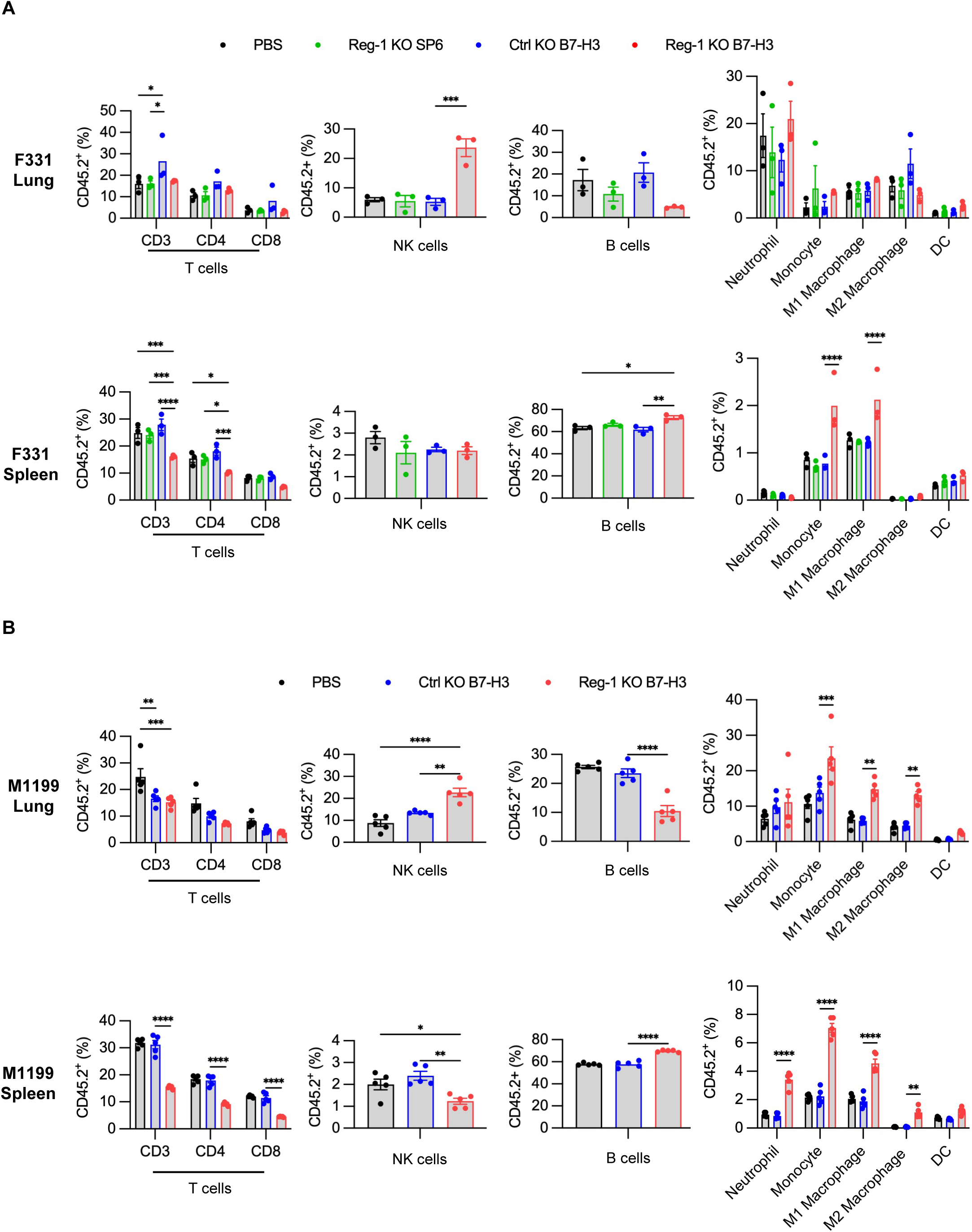
Reg-1 KO B7-H3-CAR T cells induce infiltration of NK cells. (**A, B**) Mice were injected with F331 (**A**) or M1199 (**B**) cells, followed by infusion with indicated CAR T cells as described in **Fig. S9**. (**A**) Percentages of T, NK, B and myeloid cell subsets among CD45.2^+^ viable cells in the lungs and spleens on day 7 after CAR T cell infusion (n=3 mice per group). (**B**) Percentages of T, NK, B and myeloid cell subsets among CD45.2^+^ viable cells in the lungs and spleens on day 7 after CAR T cell infusion (n=5 mice per group). NK: natural killer cell, DC: Dendritic cell. Two-way ANOVA with Tukey’s test for multiple comparisons or unpaired student’s *t*-test: *:p<0.05, **:p<0.01, ***:p<0.001, ****:p<0.0001.

**Extended data Figure 6:**
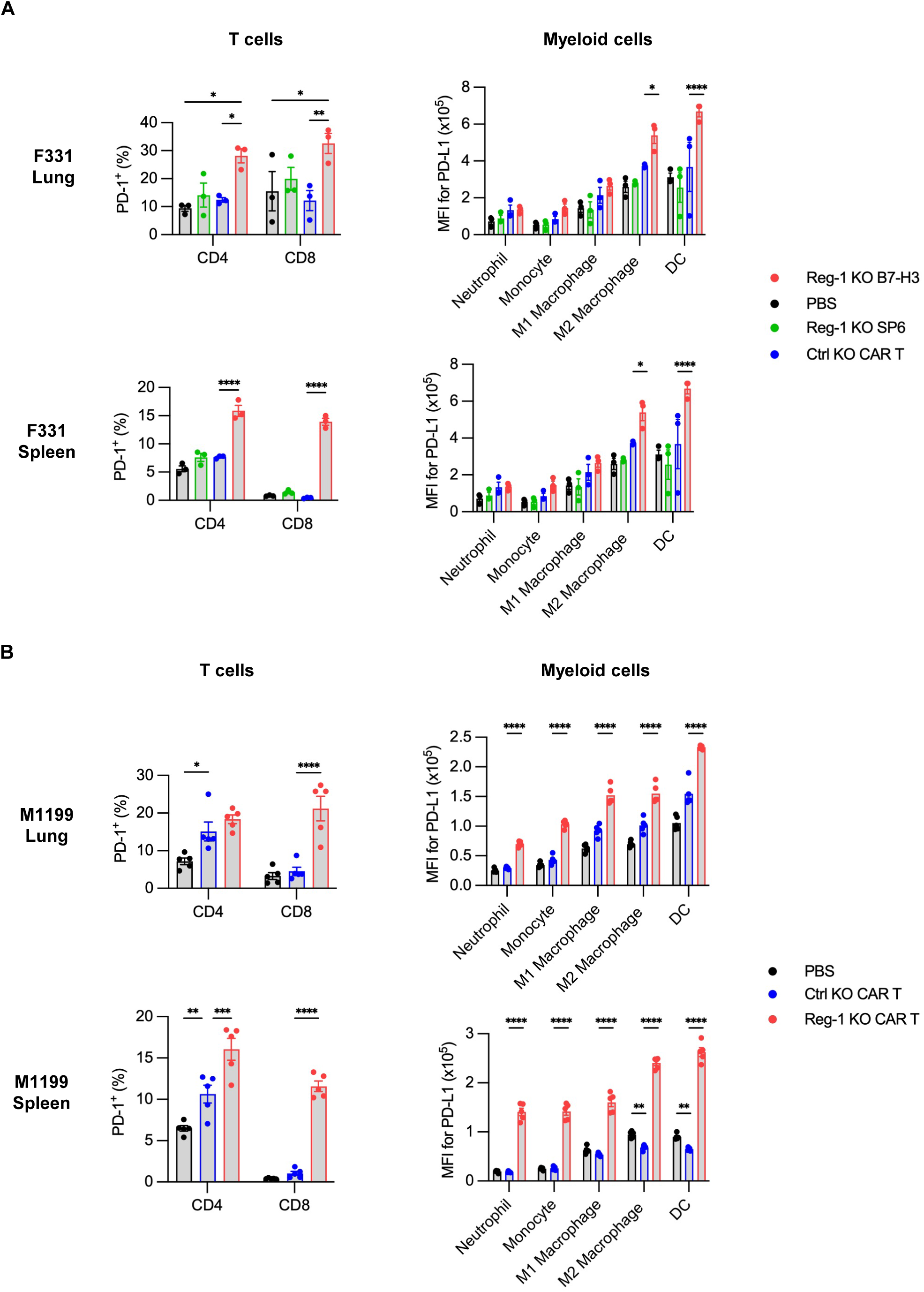
Reg-1 KO B7-H3-CAR T cells upregulate PD-1 expression on endogenous T cells and PD-L1 expression on endogenous myeloid cells. (**A, B**) Mice were injected with (**A**) F331 (n=3 mice per group) or (**B**) M1199 (n=5 mice per group) cells, followed by infusion with indicated CAR T cells as described in **Fig. S9**. Percentages of PD-1^+^ cells among endogenous CD4^+^ and CD8^+^ T cells (left) or PD-L1 expression on indicated myeloid cell populations (right) in lungs and spleens on day 7 post CAR T cell infusion, as evaluated by flow cytometry. Two-way ANOVA with Tukey’s test for multiple comparisons; *:p<0.05, **:p<0.01, ***:p<0.001, ****:p<0.0001.

**Extended data Figure 7:**
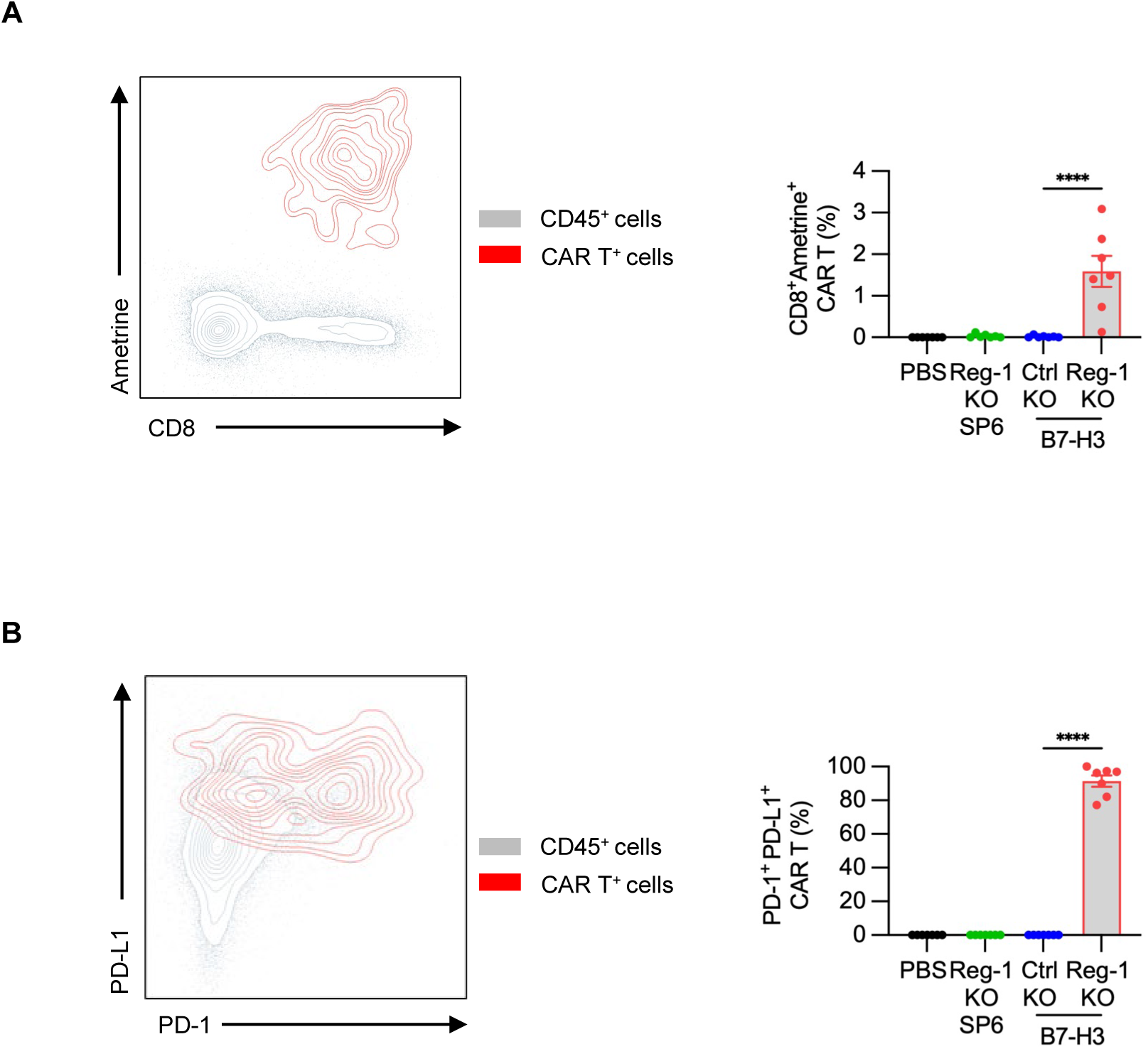
Reg-1 KO enhances B7-H3-CAR T cell persistence on day 21 post transfer. Data was pooled from all 7 mice in each condition between both immunophenotyping experiments described for Fig. 4 and **Fig. S19**. (**A**) Bar graphs gives the fraction of CAR T cells observed under each treatment condition from total CD45^+^ cells recovered, showing a unique persistence for Reg-1 KO B7-H3-CAR T cells. (**B**) Bar graphs gives the fraction of PD-1^+^PD-L1^+^ CAR T cells, indicating that nearly all CAR T cells persisting at this time point expressed these markers. Contour plots in (**A, B**) show co-expression (indicated in red) of Ametrine and CD8 (**A**) or PD-L1 and PD-1 (**B**) populations relative to concatenated data from all CD45^+^ cells (grey); Data from two independent experiments. One-way ANOVA; ****:p<0.0001;

**Extended data Figure 8:**
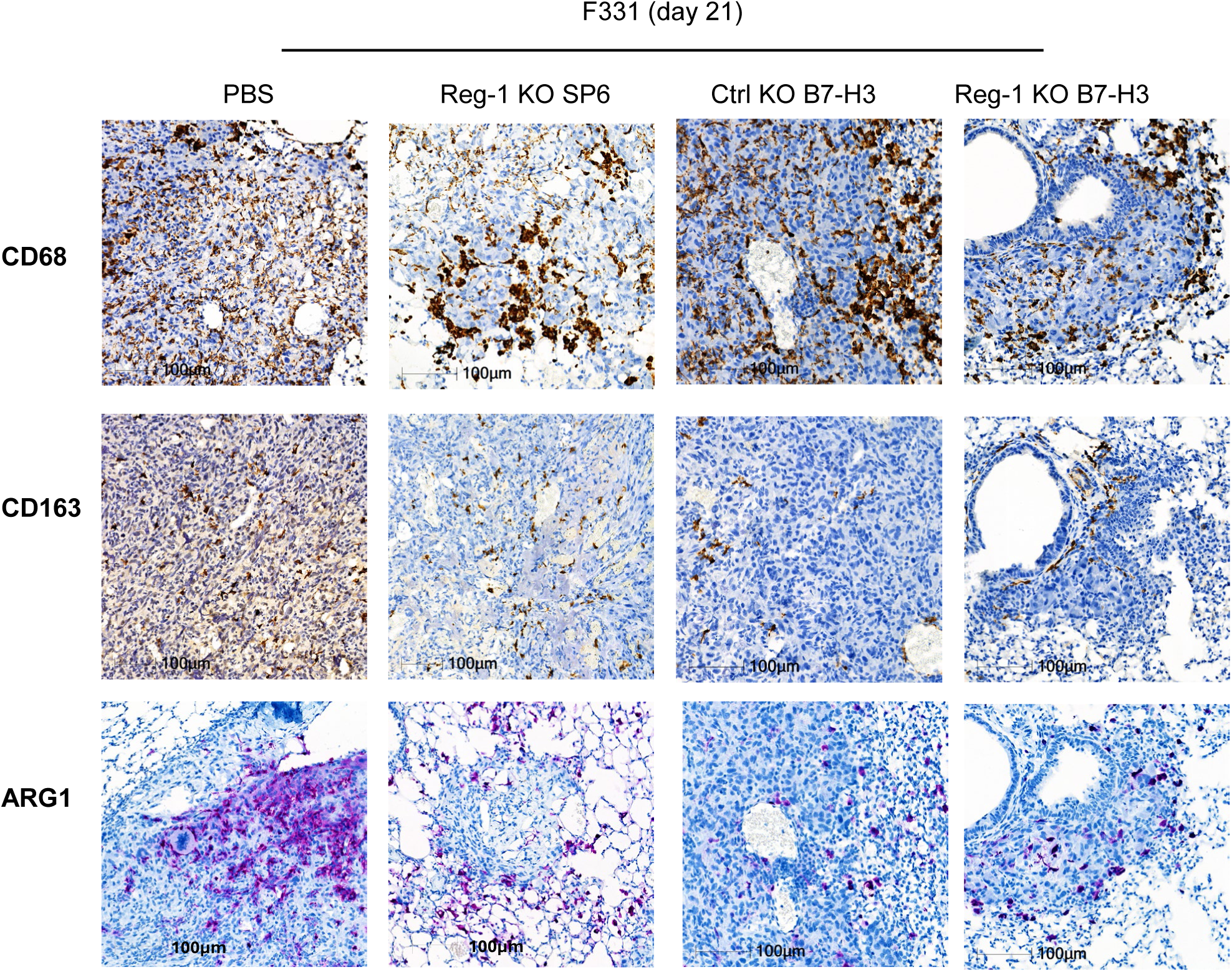
Reg-1 KO B7-H3-CAR T cells decrease macrophage signatures in osteosarcoma. IHC for a universal-macrophage marker (CD68 staining) or specific markers for M2 macrophages (CD163 and ARG1) in lung tumors from F331 tumor-bearing mice on day 21 after infusion with Reg-1 KO SP6, Ctrl KO or Reg-1 KO B7-H3-CAR T cells. 3,3’-Diaminobenzidine (DAB) chromogen with hematoxylin counterstain. Representative images. 20× magnification, 100 µm scale.

**Extended data Figure 9:**
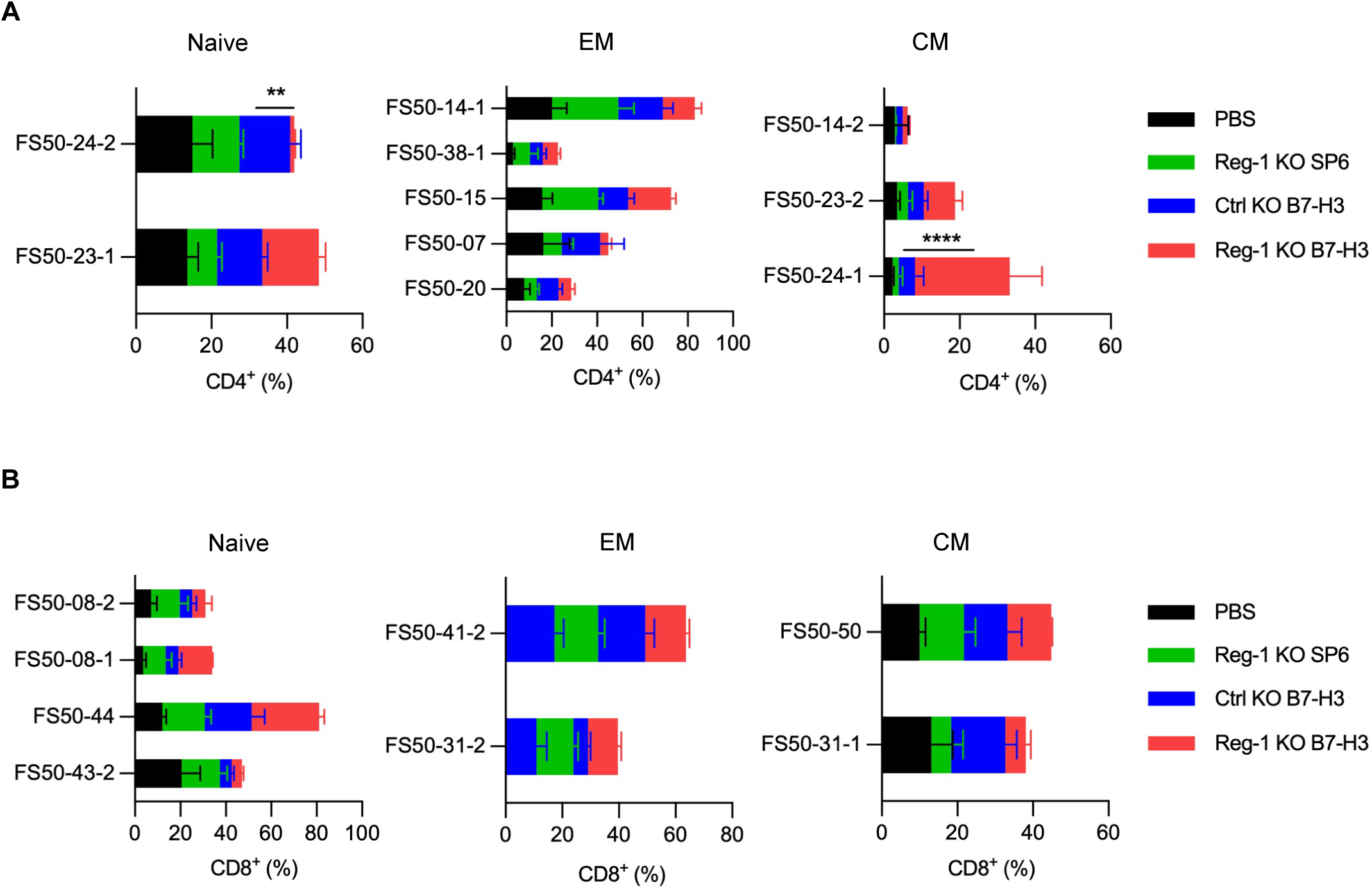
Reg-1 KO B7-H3-CAR T cells alter endogenous T cell subsets. Cluster distribution of the experiment described in Fig. 4 and Fig. 5G**,H**. (**A, B**) The naive, EM and CM subsets of (**A**) CD4^+^ and (**B**) CD8^+^ T cell clusters from the experiment reported in Fig. 5**. G,H** and illustrated in**Table S2** (n=3 mice per group). Comparison was performed for the Reg-1 KO B7-H3-CAR T treatment condition relative to the Ctrl KO B7-H3-CAR T cell recipients; two-way ANOVA, Tukey’s multiple comparisons test; **:p<0.01; ****:p<0.0001.

**Extended data Figure 10:**
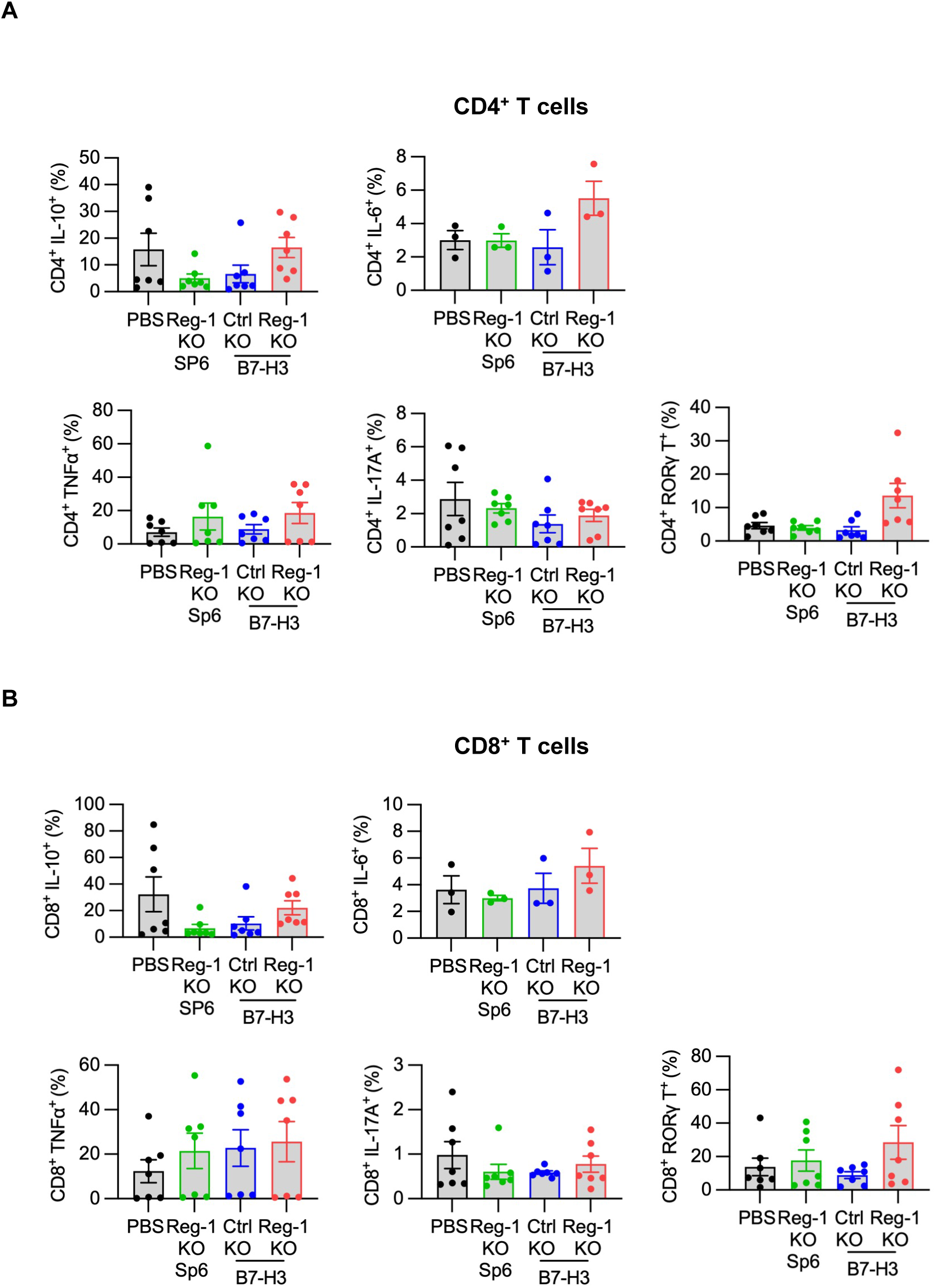
Cytokine production of endogenous T cells post-CAR T cell infusion. Intracellular flow cytometry panel detailed in **Table S5** and outlined in methods was conducted on an aliquot of single-cell suspensions from lungs of F331 tumor-bearing mice that received infusion of the indicated CAR T cells, as described in Fig. 4 and **Fig. S19**. (**A,B**) Percentages of IL-10, IL-6, TNFα, IL-17A, and RORγ-expressing cells among (**A**) CD4^+^ and (**B**) CD8^+^ T cells.

## Supplementary Figures

**Figure S1:** Generation of Reg-1 KO B7-H3-CAR T cells

**Figure S2:** B7-H3 expression on OS cell lines

**Figure S3:** Reg-1 KO B7-H3-CAR T cells demonstrate antigen specific killing *in vitro*

**Figure S4:** Changes in body weight post-CAR T cell infusion

**Figure S5:** Osteosarcomas continue to express B7-H3 post-CAR T cell infusion

**Figure S6:** Reg-1 KO did not alter PD-1 expression on B7-H3-CAR T cells

**Figure S7:** Endogenous CD4^+^ and CD8^+^ T cell differentiation post-CAR T cell infusion

**Figure S8:** PD-L1 blockade does not improve Reg-1 KO B7-H3-CAR T cells antitumor activity

**Figure S9:** Reg-1 KO increased the frequency of clonally diverse CAR T cells in the lung

**Figure S10:** t-SNE association and Flow-SOM clustering of high dimensional flow cytometry analysis plots

**Figure S11:** Strategy for phenotyping FlowSOM clusters of lung isolates post-CAR T cell infusion

**Figure S12:** Marker expression across t-SNE associations in high-dimensional phenotyping analysis

**Figure S13:** Frequency of distinct immune cells subsets among CD45^+^ cells

**Figure S14:** Reg-1 KO B7-H3-CAR T cells induced changes in lung infiltrating fibroblasts

## Supplementary Tables

**Table S1:** List of extracellular flow cytometry panel with 23 fluorophores

**Table S2:** List of FlowSOM clusters from 23 fluorophore flow cytometry panel

**Table S3:** List of extracellular flow cytometry panel with 38 fluorophores

**Table S4:** List of FlowSOM clusters from 38 fluorophore flow cytometry panel

**Table S5:** List of intracellular flow cytometry panel with 25 fluorophores

**Table S6:** Antibodies and methods used for immunohistochemistry

