## Supplementary Figures 1 to 14 for "Targeting Regnase-1 unleashes CAR T cell antitumor activity for osteosarcoma and creates a proinflammatory tumor microenvironment"

[1]Department of Bone Marrow Transplantation and Cellular Therapy, St. Jude Children's Research Hospital, Memphis, TN; [2]Department of Immunology, St. Jude Children's Research Hospital, Memphis, TN; [3]Department of Pathology, St. Jude Children's Research Hospital, Memphis, TN; [4]Flow Cytometry and Cell Sorting Shared Resource, St. Jude Children's Research Hospital, Memphis, TN; [5]Center for Advanced Genome Engineering, St. Jude Children's Research Hospital, Memphis, TN; [6]Aflac Cancer and Blood Disorders Center and the Winship Cancer Institute, Emory University, Atlanta, GA

[#]These authors share senior authorship

**Corresponding Authors**

Stephen Gottschalk, Department of Bone Marrow Transplantation and Cellular Therapy, St. Jude Children's Research Hospital, 262 Danny Thomas Place, Memphis, TN 38105, Phone: (901)-595-5935,

Hongbo Chi, Department of Immunology, St. Jude Children's Research Hospital, 262 Danny Thomas Place, Memphis, TN 38105, Phone: (901)-595-6282,

**Supplementary Figure 1**

**A**

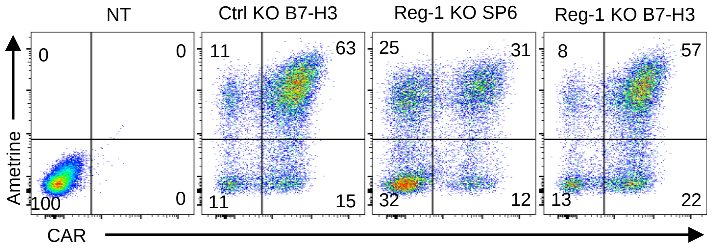


**B**

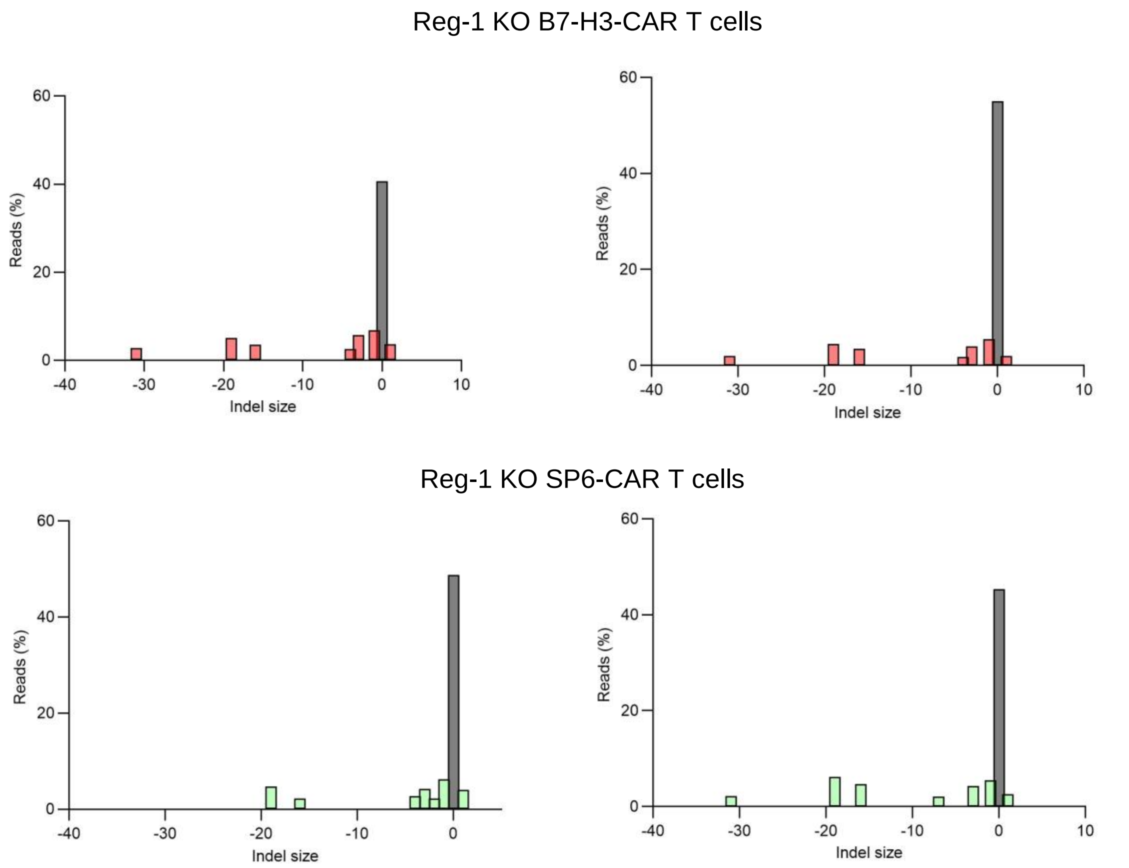


**C**

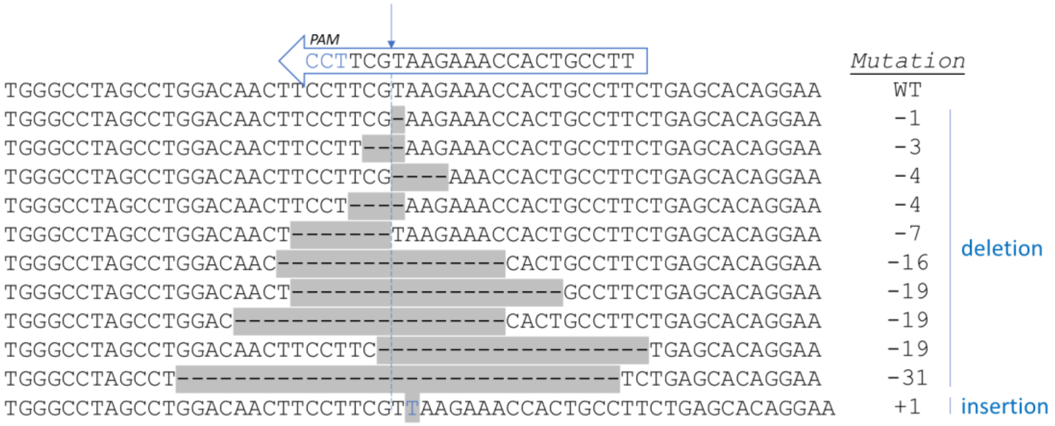


*Regnase1 sgRNA-1*

5’-AAGGCAGTGGTTTCTTACGA-3’

**Figure S1: Generation of Reg-1 KO B7-H3-CAR T cells**

(**A**) Representative flow cytometry plots to evaluate co-transduction efficiency of indicated sgRNA (Ametrine$^{+}$) and CAR expression in NT, Ctrl KO B7-H3 CAR, Reg-1 KO SP6-CAR, or Reg-1 KO B7-H3-CAR T cells on day 3 post-transduction. (**B**) Top 8 insertion and deletion (Indel) sites in Reg-1 KO SP6 or B7-H3 CAR T cells determined via deep sequencing analysis at the exonic target site of the Reg-1 sgRNA. Top 8 indels compromised 71.1% and 59.3% in Reg-1 KO B7-H3-CAR T cells and 62.3% and 51.2% in controls. Data from 2 independent experiments is shown. (**C**) Indel sequencing results in Reg-1 KO B7-H3-CAR T cells.

**Supplementary Figure 2**

**A**

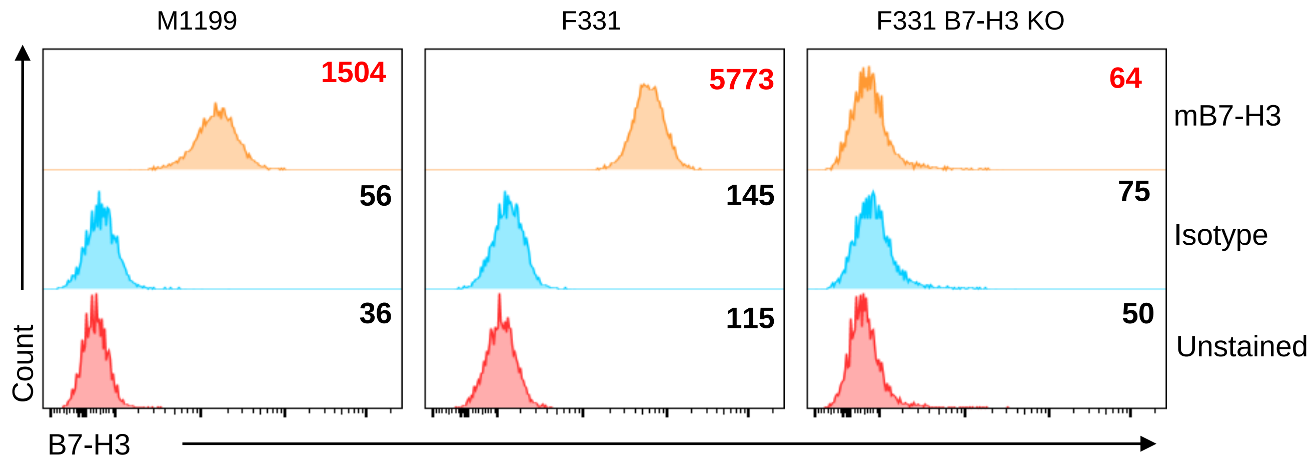


**B**

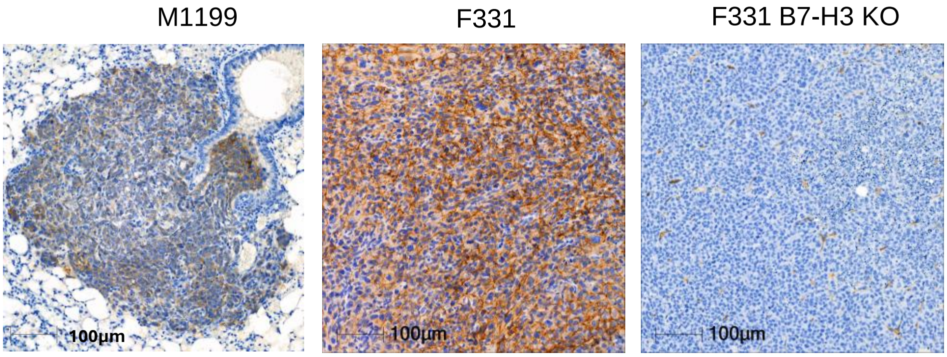

**Figure S2: B7-H3 expression on OS cell lines**

(**A**) Representative flow cytometry plot showing B7-H3 expression on M1199 and F331; F331 B7-H3 KO OS cell lines was included as a negative control; Red = Unstained, Blue = Isotype control, Orange = B7-H3 staining. Indicated values on upper right corners represents mean fluorescence intensity (MFI). (**B**) B7-H3 expression was examined by IHC for lung tumors; representative images on day 28 post M1199 or F331 i.v. injection. F331 B7-H3 KO cells were included as a negative control. 3,3'-Diaminobenzidine (DAB) chromogen with hematoxylin counterstain. 20x magnification, 100 µm scale.

### Supplementary Figure 3

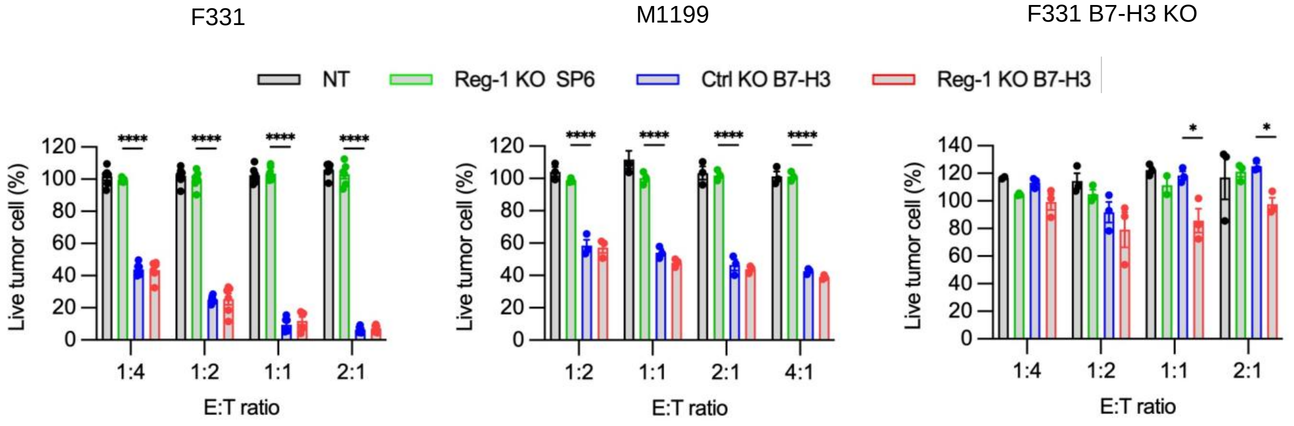

**Figure S3: Reg-1 KO B7-H3-CAR T cells demonstrate antigen specific killing *in vitro***

3-(4,5-dimethylthiazol-2-yl)-5-(3-carboxymethoxyphenyl)-2-(4-sulfophenyl)-2H-tetrazolium (MTS) assay with B7-H3-positive (F331, M1199) and B7-H3-negative (F331 B7-H3 KO) cell lines at indicated effector to target (E:T) ratios; n=3, two-way ANOVA with Tukey's test for multiple comparisons; *:$p<0.05$, ****:$p<0.0001$.

#### Supplementary Figure 4

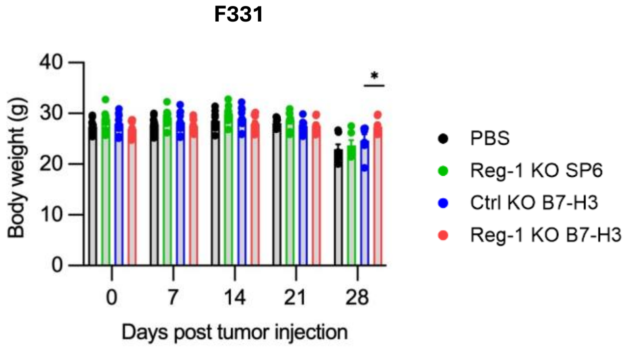


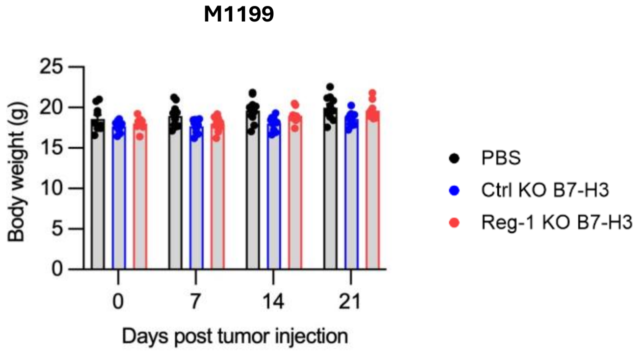

**Figure S4: Changes in body weight post-CAR T cell infusion**

Weekly body weight measurement of F331 and M1199 bearing mice post infusion with Reg-1 KO SP6, Ctrl KO or Reg-1 KO B7-H3-CAR T cells. PBS served as the control. Two-way ANOVA with Tukey's test for multiple comparisons; *:$p<0.05$.

#### Supplementary Figure 5

A

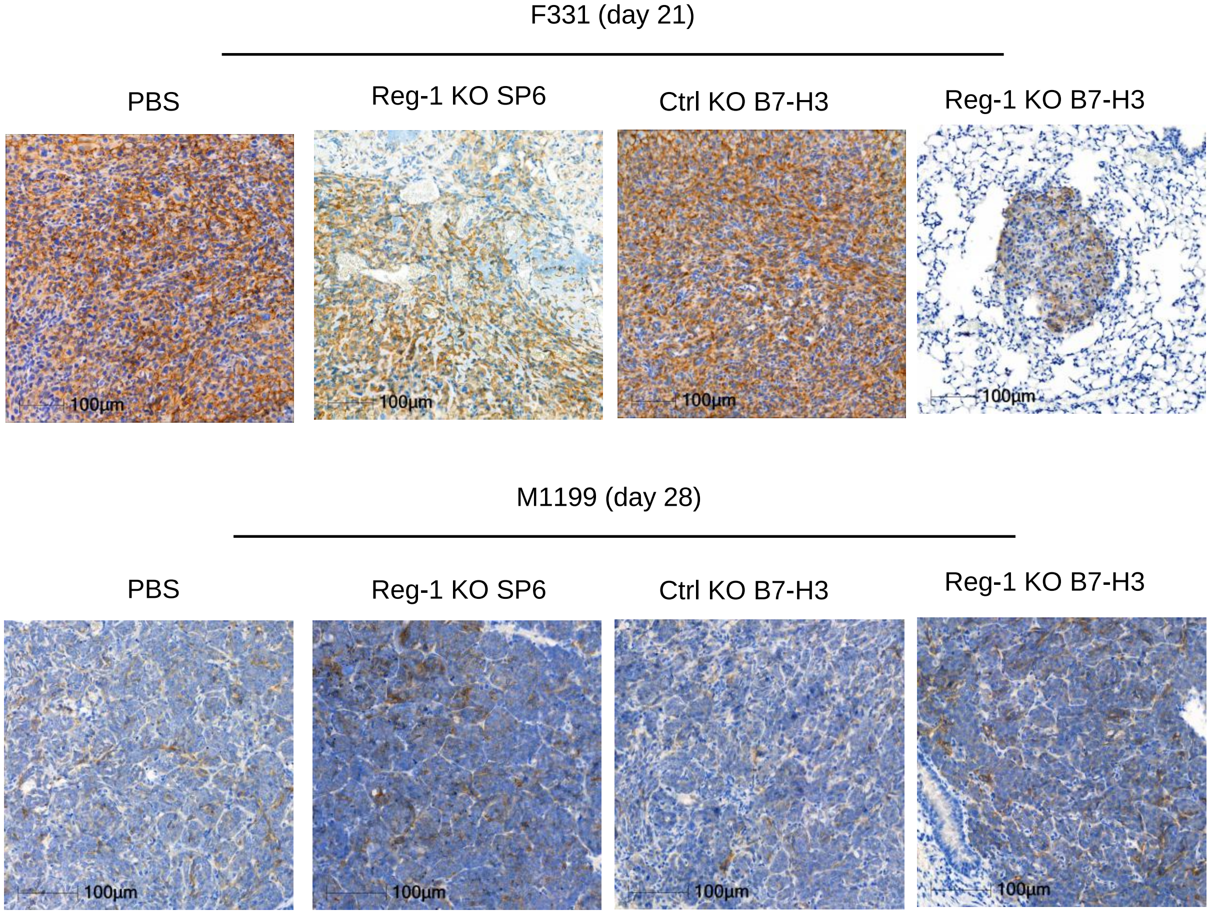


B

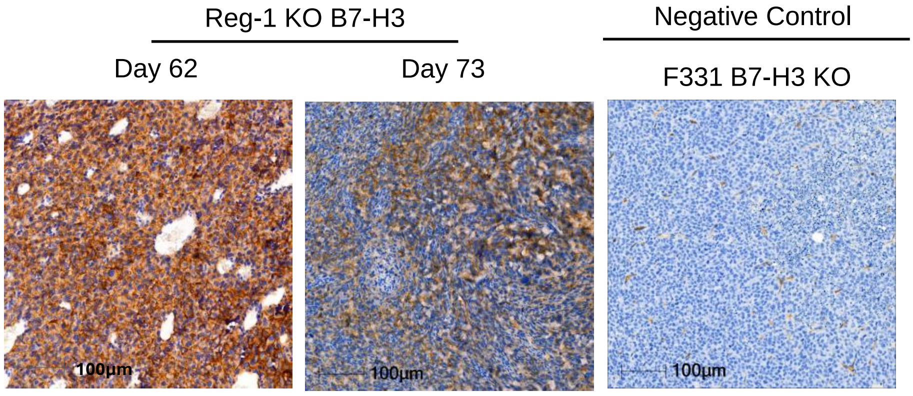

**Figure S5: Osteosarcomas continue to express B7-H3 post-CAR T cell infusion**
(**A**) IHC for B7-H3 expression in lung tumors from F331 and M1199 bearing mice on day 21 (F331; upper) or 28 (M1199; lower) after infusion with Reg-1 KO SP6, Ctrl KO or Reg-1 KO B7-H3-CAR T cells. (**B**) IHC for B7-H3 expression in recurring lungs F331 tumors on days 62 and 73 after Reg-KO B7-H3-CAR T cell infusion. F331 B7-H3 KO lung tumors served as a negative control. 3,3'-Diaminobenzidine (DAB) chromogen with hematoxylin counterstain. 20x magnification, 100 μm scale.

#### Supplementary Figure 6

A

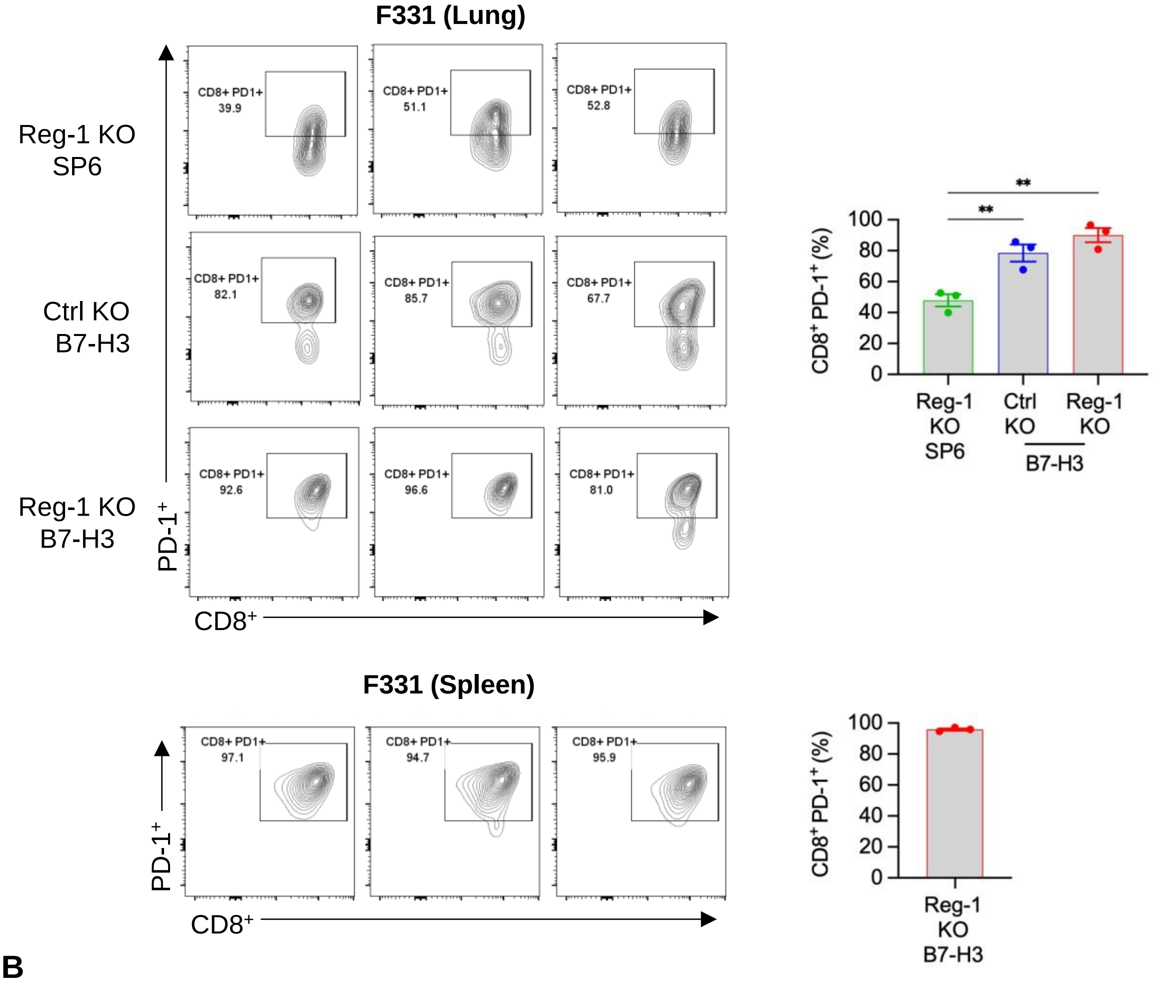


B

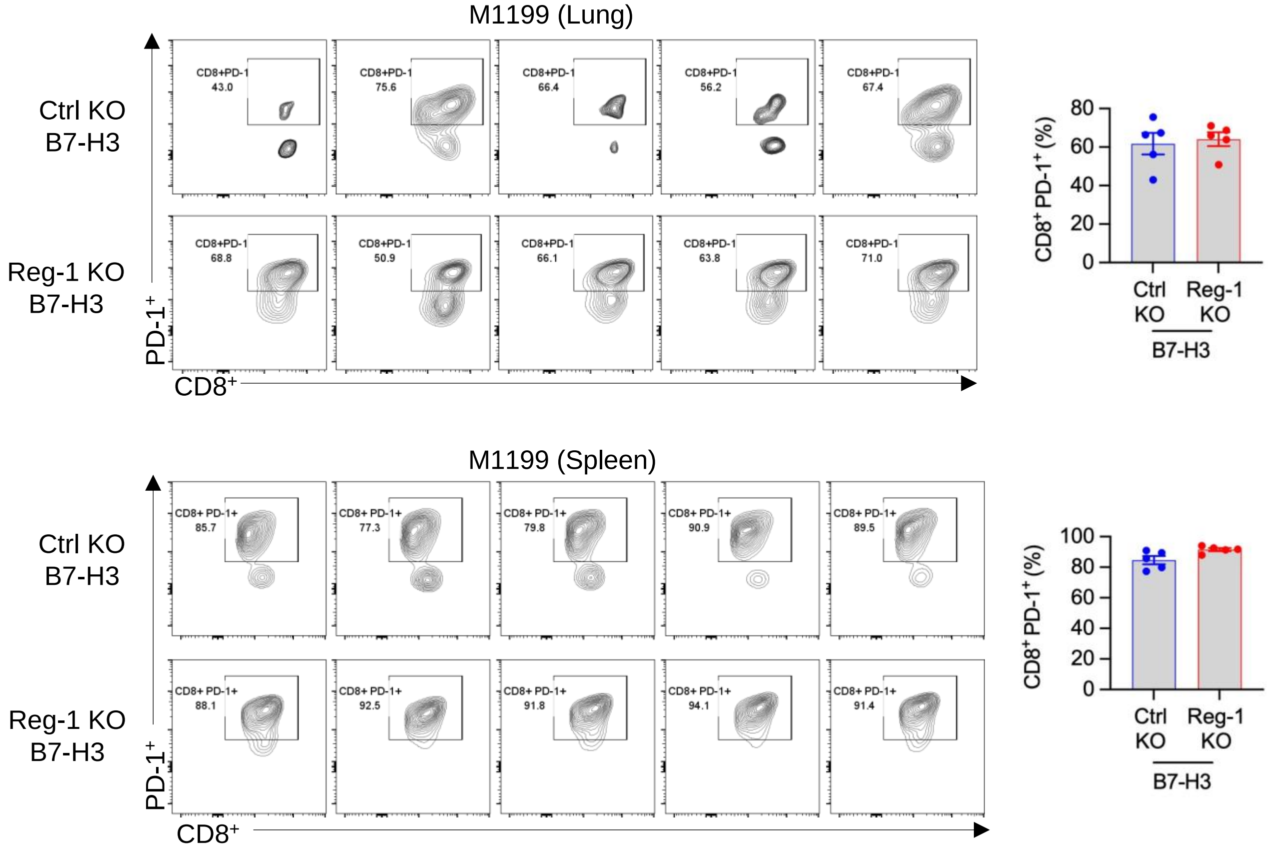

**Figure S6: Reg-1 KO did not alter PD-1 expression on B7-H3-CAR T cells**

(**A**) C57BL/6 mice were injected with $1x10^6$ F331 (i.v.) and on day 7 received a single i.v. dose of $1x10^6$ Reg-1 KO SP6, Ctrl KO or Reg-1 KO B7-H3-CAR T cells. PBS-treated mice served as control. Flow cytometry plots and quantification of PD-1 expression on $CD8^+$ CAR T cells in the lungs and spleens on day 7 post CAR T cell infusion; n=3 mice per group (there are not enough CAR T cells in the spleens of Reg-1 KO SP6 and Ctrl KO B7-H3-CAR T cell-treated mice to make a comparison). (**B**) C57BL/6 mice were implanted with $2x10^5$ M1199 (i.v.) and on day 7 received a single i.v. dose of $2x10^6$ Ctrl KO or Reg-1 KO B7-H3-CAR T cells. PBS-treated mice served as control. Flow cytometry plots and quantification of PD-1 expression on $CD8^+$ B7-H3-CAR T cells in the lungs and spleens on day 7 post CAR T cell infusion, n=5 mice per group. One-way ANOVA, **:p<0.01, Turkey's multiple comparison test.

#### Supplementary Figure 7

A

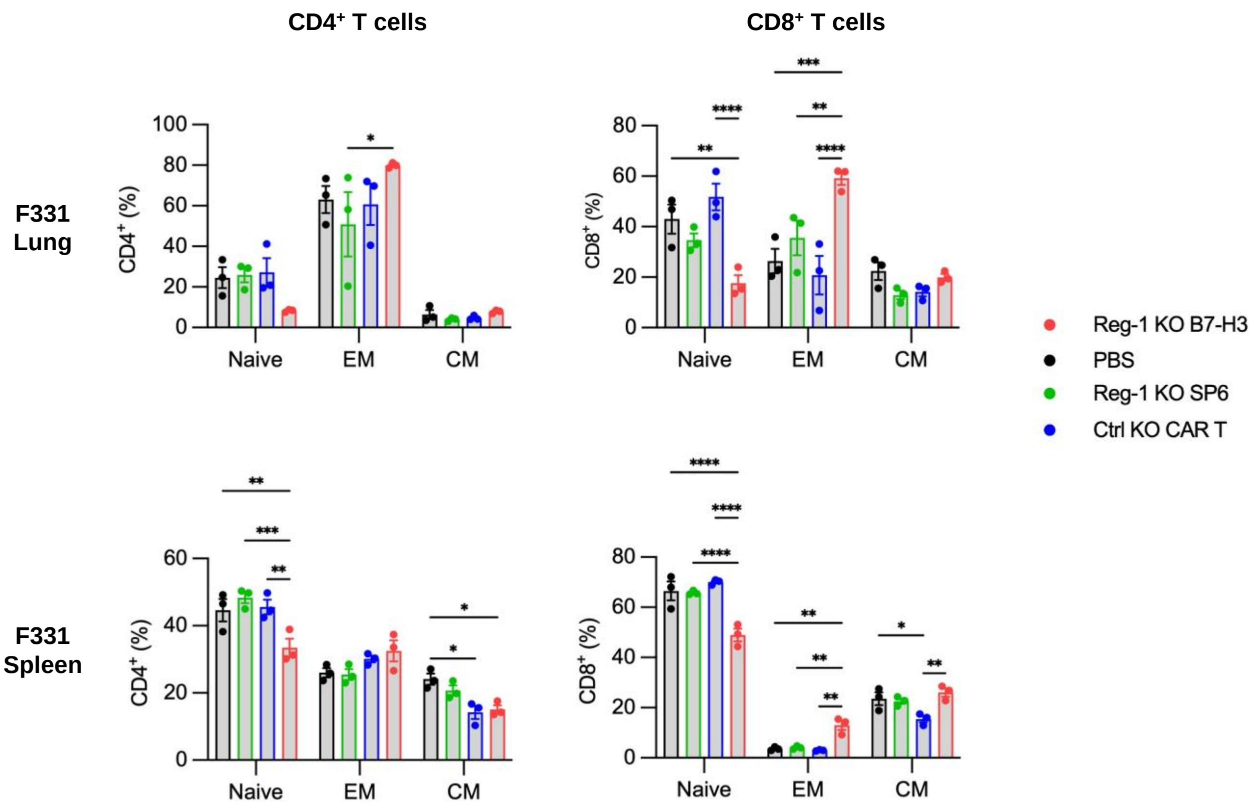


B

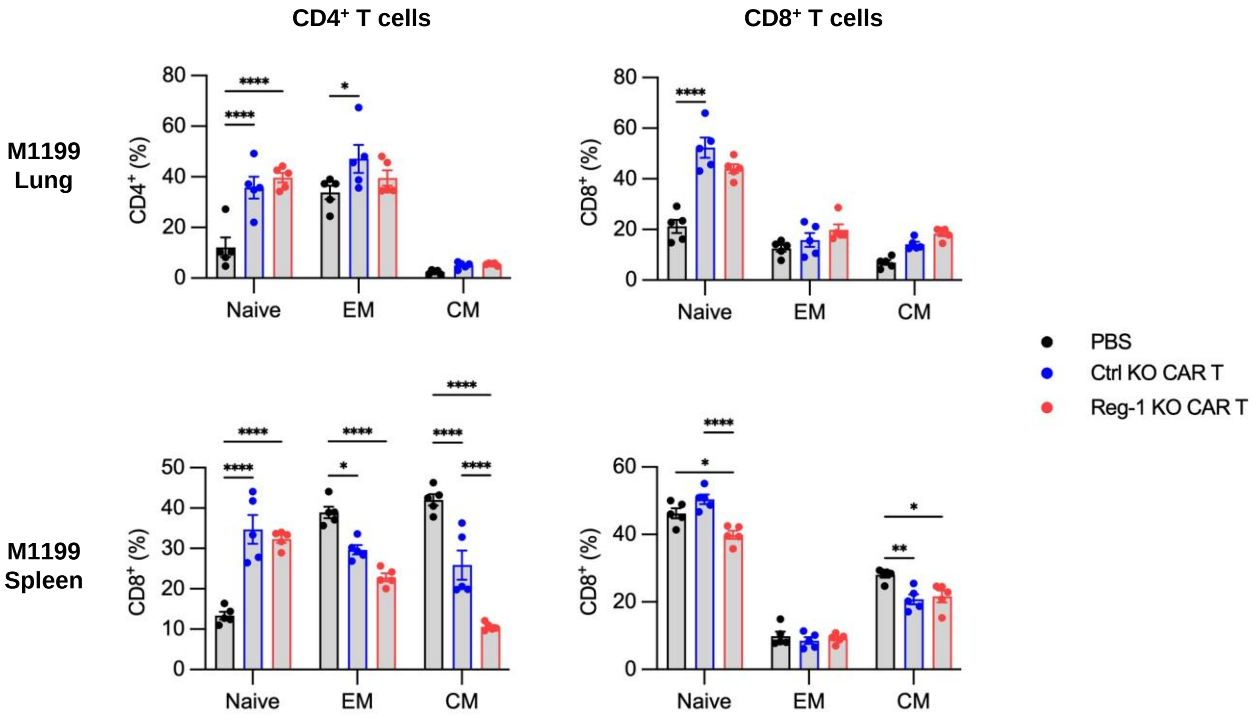

**Figure S7: Endogenous CD4$^+$ and CD8$^+$ T cell differentiation post-CAR T cell infusion**
(**A, B**) Mice were injected with (**A**) F331 (n=3 mice per group) or (**B**) M1199 (n=5 mice per group) cells, followed by infusion with indicated CAR T cells as described in **Fig. S9**. Percentages of endogenous naïve (CD62L$^+$CD44$^-$), effector memory (EM; CD62L$^-$CD44$^+$) and central memory (CM; CD62L$^+$CD44$^+$) populations among CD4$^+$ (left) and CD8$^+$ (right) T cells in lungs and spleens of mice on day 7 post CAR T cell infusion, as evaluated by flow cytometry. Two-way ANOVA with Tukey's test for multiple comparisons; *:$p<0.05$, **:$p<0.01$, ***:$p<0.001$, ****:$p<0.0001$.

#### Supplementary Figure 8

**A**

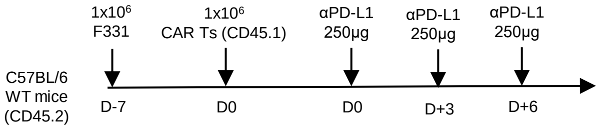


**B**

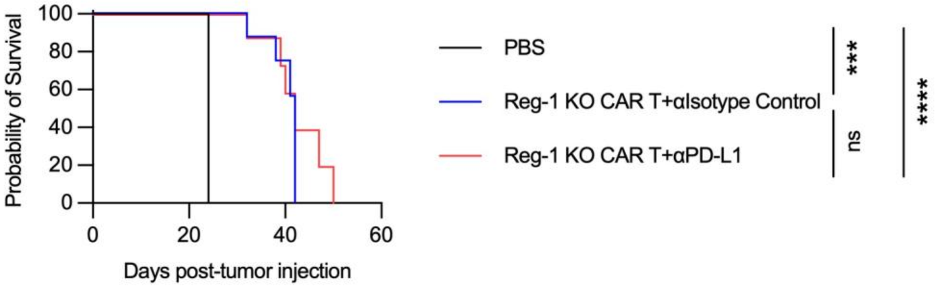

**Figure S8: PD-L1 blockade does not improve Reg-1 KO B7-H3-CAR T cells antitumor activity**

(**A**) Experimental schematic. (**B**) Kaplan-Meier survival curve for F331 tumor-bearing mice that received Reg-KO B7-H3-CAR T cells with or without PD-L1 antibody as indicated (n=5 mice per group). Log-rank Mantel-Cox test; ****: p<0.0001.

#### Supplementary Figure 9

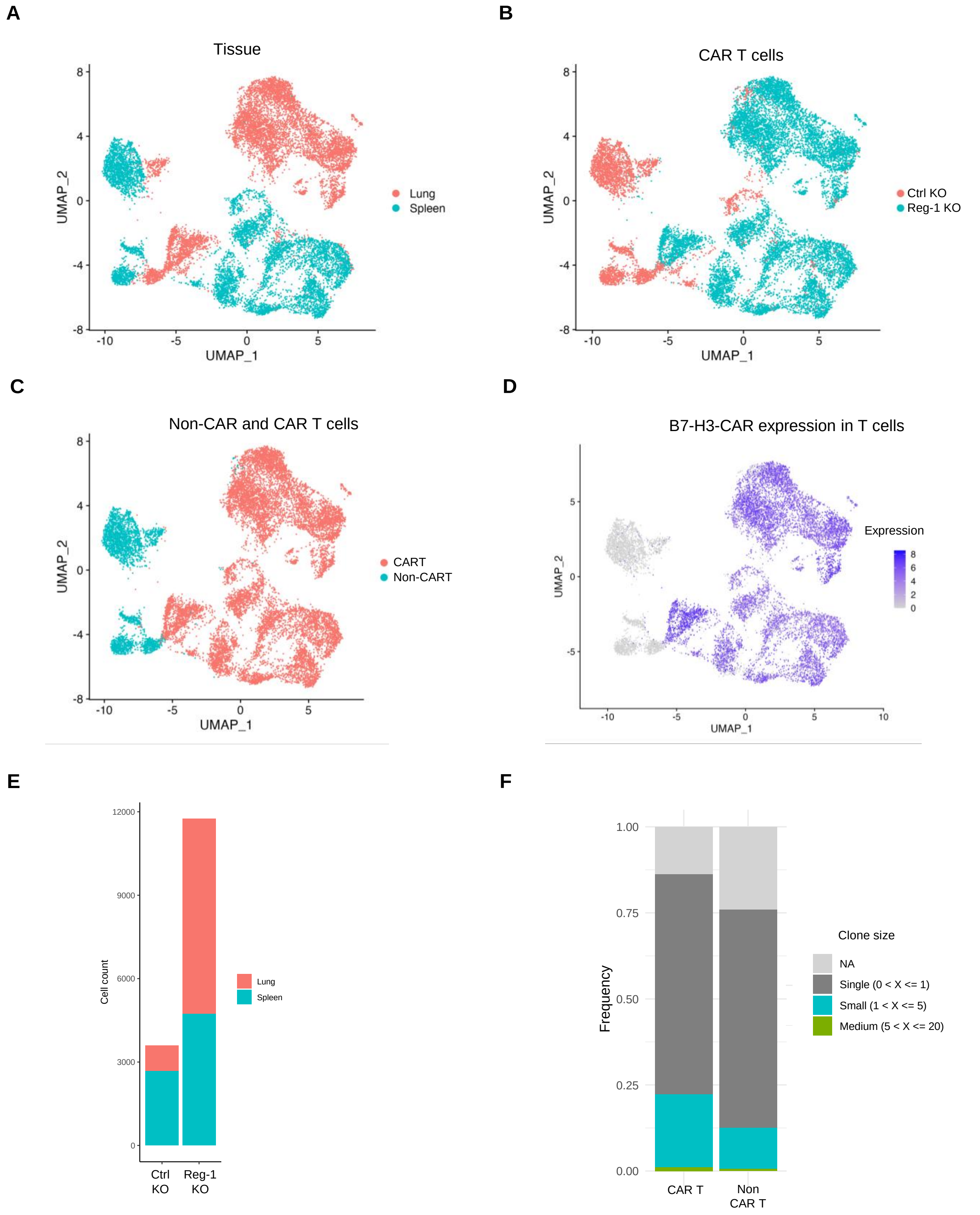

**Figure S9: Reg-1 KO increased the frequency of clonally diverse CAR T cells in the lung**
C57BL/6 mice ($CD45.2^+$) received F331 cells (i.v.) followed by a single dose of $5x10^6$ Reg-1 KO or Ctrl KO B7-H3- CAR T cells ($CD45.1^+$) on day 7 post tumor cell injection. Lungs and spleens were isolated on day 5 post CAR T cell infusion, and cells were pooled from 3 mice for scRNA-seq profiling. (**A–D**) UMAP projection of (**A**) B7-H3-CAR T cells in the lungs and spleens, (**B**) Reg-1 KO and Ctrl KO B7-H3-CAR T cells within the lungs and spleens, (**C,D**) UMAP distrbution of B7-H3-CAR T and non-CAR T cells and (**D**) B7-H3-CAR expression in the lungs and spleens. (**E**) Cell number of Reg-1 KO compared to Ctrl KO B7-H3-CAR T cells in lungs and spleens. (**F**) scTCR-seq analysis showing T cell clonal diversity in non-CAR and B7-H3-CAR T cells.

**Supplementary Figure 10**

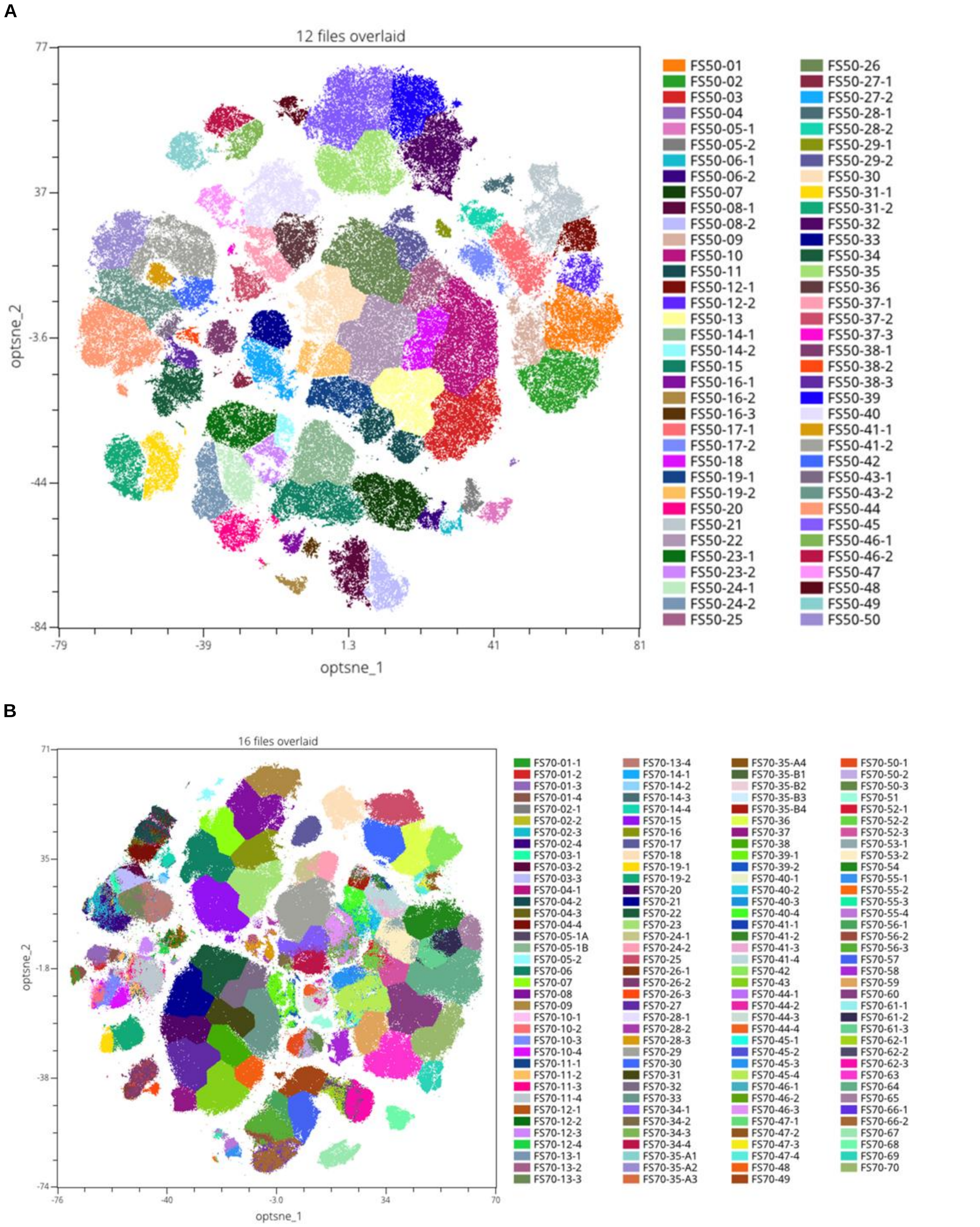

**Figure S10: t-SNE association and Flow-SOM clustering of high dimensional flow cytometry analysis plots**

C57BL/6 mice were injected with $1x10^6$ F331 (i.v.) and on day 7 received a single i.v. dose of $1x10^6$ Reg-1 KO SP6, Ctrl KO or Reg-1 KO B7-H3-CAR T cells. PBS treated mice served as control. On day 21 post CAR T cell infusion, mice were euthanized and lungs isolated for high dimensional flow cytometry analysis. Unsupervised clustering analysis was performed using the OMIQ analysis software suite by application of the FlowSOM algorithm based on t-SNE marker associations, as described for **Fig 4**. (**A**) High dimensional reduction plot from a 23-color panel concatenated from 12 samples, resulting in 72 distinct clusters and subclusters. Marker expression details are listed in **Table S2**, and the schematic used for phenotypic classification is outlined in **Fig. S17** (n=3 mice per group, and 26,000 viable $CD45^+$ cells analyzed per mouse). (**B**) High dimensional reduction plot from the second experiment, this time employing a 38-color panel. Concatenated results from 16 samples resulting in 147 distinct clusters and subclusters are shown. Marker expression details are listed in **Table S4**, and phenotypic classification followed the same outline as shown in **Fig. S17** (n=4 mice per group, and 72,000 viable $CD45^+$ cells were analyzed per mouse). t-SNE, t-distributed stochastic neighbor embedding.

### Supplementary Figure 11

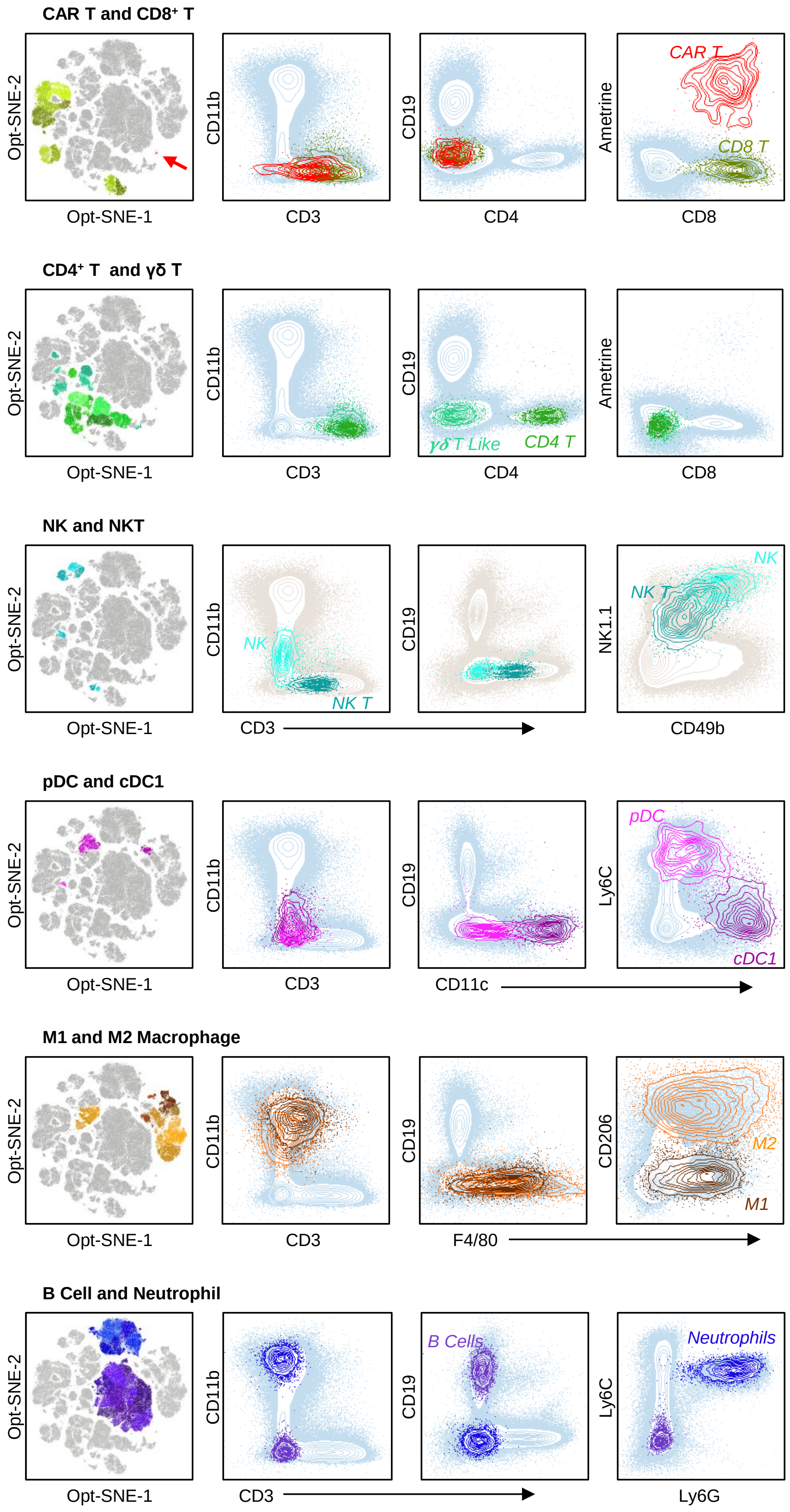

**Figure S11: Strategy for phenotyping FlowSOM clusters of lungs isolate post-CAR T cell infusion**

Clusters in both experiments described in **Fig. S16** were each phenotypically identified as illustrated here. Specifically, colored clusters from the t-SNE plot (far left panels) were assignments based on the cell surface expression markers designated in the contour plots, moving progressively from left to right. Contour plots show one representative cluster in each general assignment category from **Fig. 4A** and **Fig. S19A**. Further details of each cluster are given in **Tables S2** and **S4**. Red arrow on the top left panel highlights the small cluster of CAR T cells found in the lungs after infusion with Reg-1 KO B7-H3-CAR T cells. t-SNE, t-distributed stochastic neighbor embedding.

**Supplementary Figure 12**

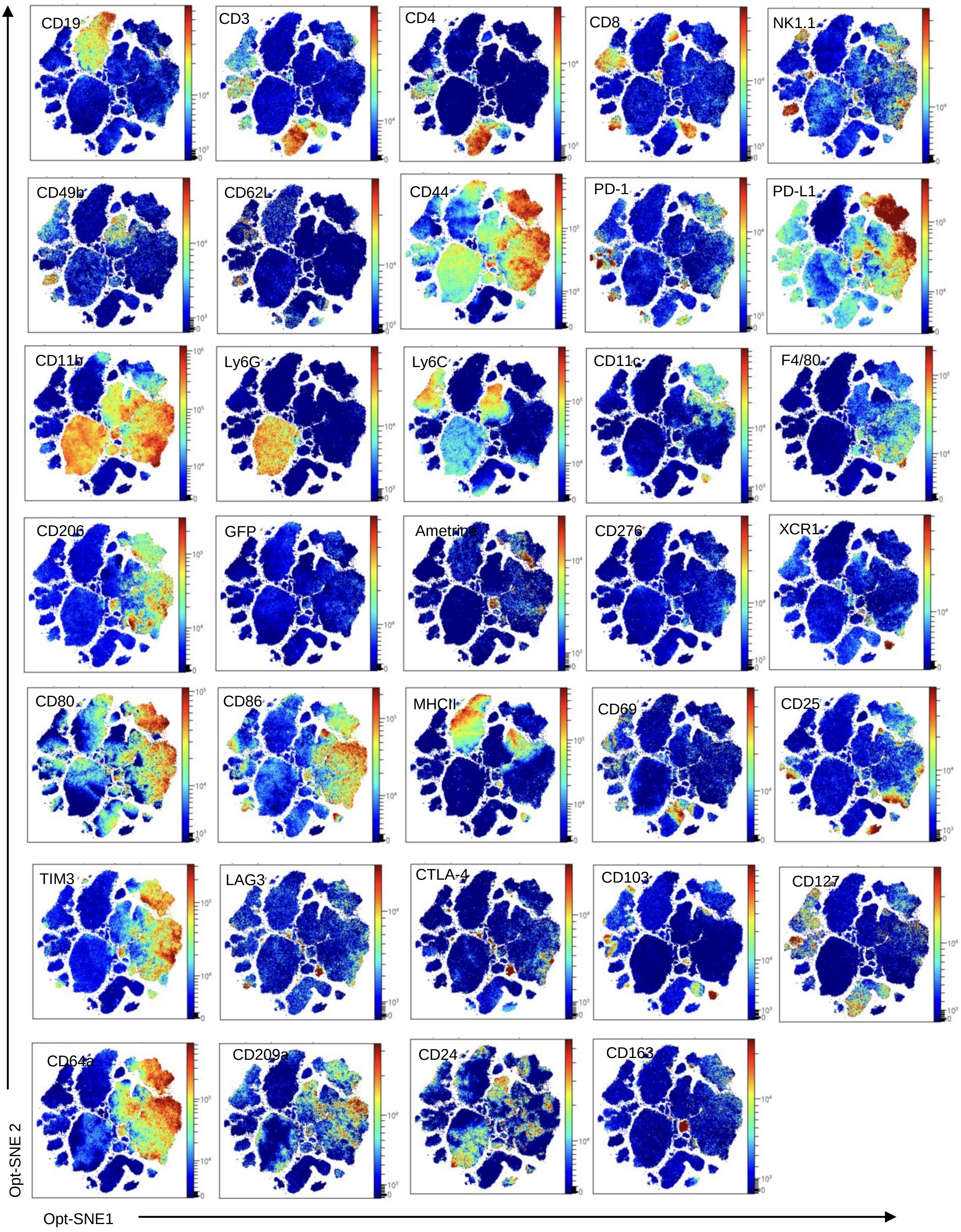

**Figure S12: Marker expression across t-SNE associations in high-dimensional phenotyping analysis**

Concatenated data from all 16 mice analyzed with the 38-marker panel described for **Fig. S16**. Heatmaps present the relative mean fluorescence intensity of the immune markers assayed, localized across the several t-SNE clusters. t-SNE, t-distributed stochastic neighbor embedding.

### Supplementary Figure 13

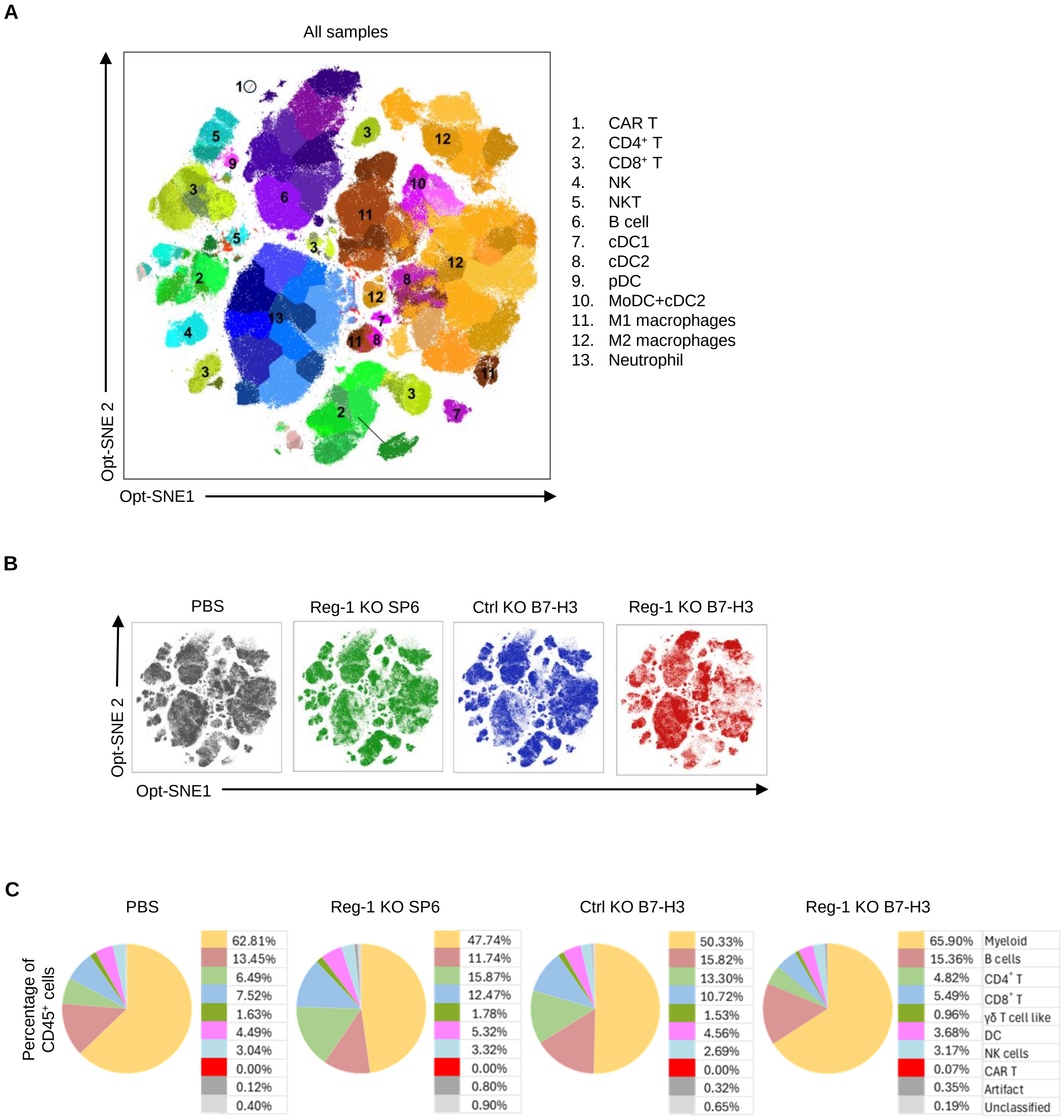

**Figure S13: Frequency of distinct immune cells subsets among CD45$^+$ cells**

(**A**) t-SNE analysis of concatenated data from all 16 mice analyzed with the 38-marker panel described for **Fig. S16** with phenotyping conducted as described for **Fig. S17**. (**B**) t-SNE plots for individual treatment conditions (n=4 mice per group) displaying the contributions of each test group to the concatenated data given in (A), and illustrating distinct cluster aggregations between groups. (**C**) Fraction of CD45$^+$ cells assigned to each immune cell subset, given as the mean from the 4 individuals in the indicated treatment group. t-SNE, t-distributed stochastic neighbor embedding.

**Supplementary Figure 14**

**A**

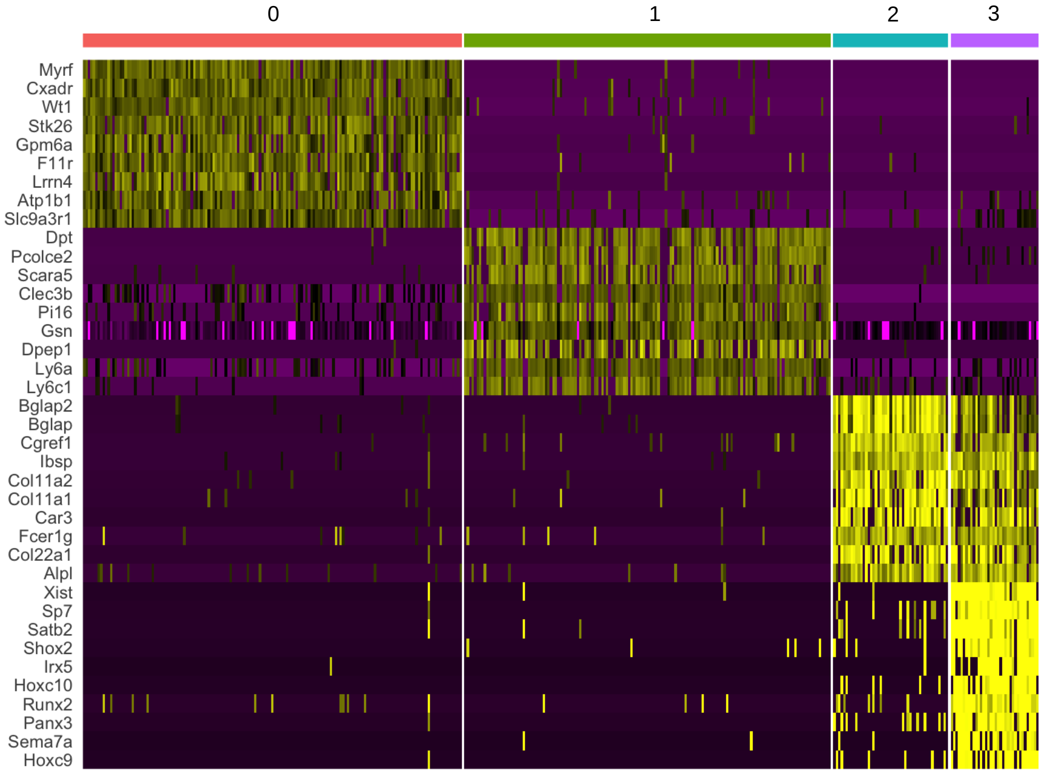


**B**

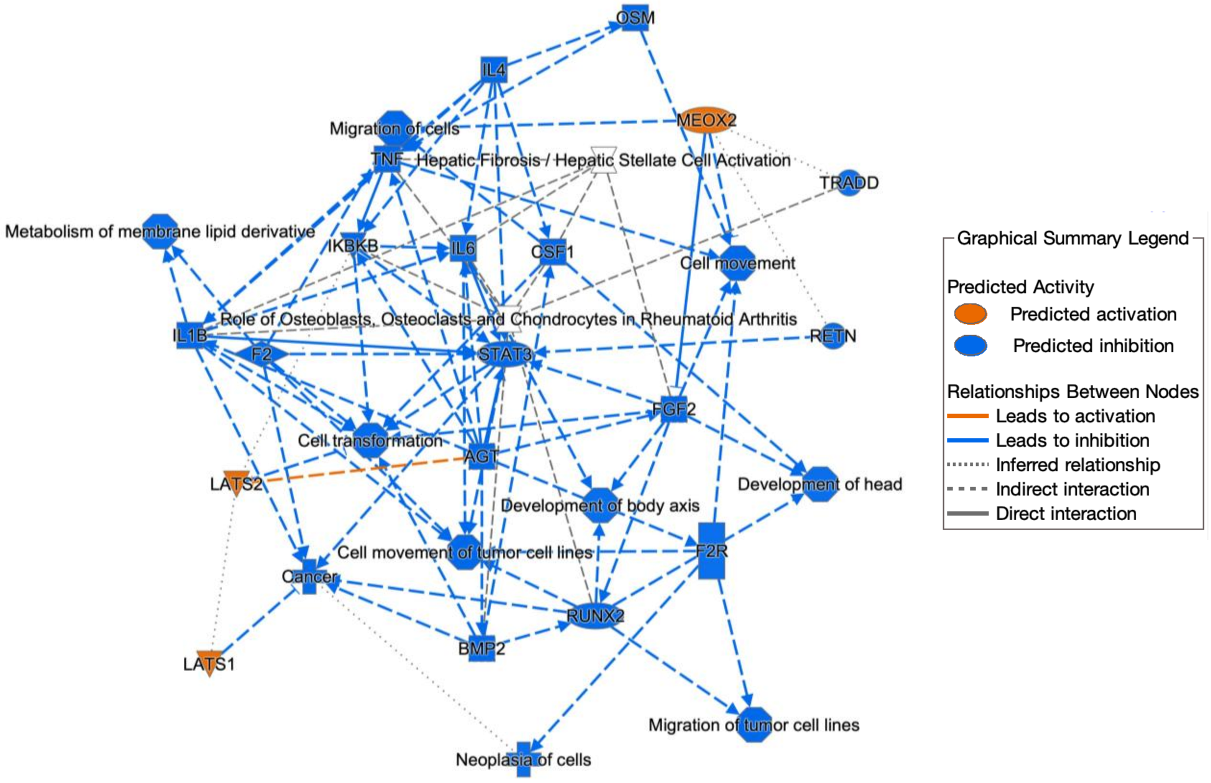

**Figure S14: Reg-1 KO B7-H3-CAR T cells induced changes in lungs infiltrating fibroblasts**
See **Fig. S15** for description of animal experiment. (**A**) Heatmap reveals differentially expressed genes in Reg-1 KO B7-H3-CAR T cells in the lungs compared to Ctrl KO B7-H3-CAR T cells in four distinct fibroblast clusters. (**B**) Ingenuity pathway analysis of fibroblasts in the lungs of mice that received Reg-1 KO compared to Ctrl KO B7-H3-CAR T cells.
